# Land-use intensity overrides grazing and precipitation effects on soil microbial communities in a subtropical agroecosystem

**DOI:** 10.64898/2026.04.30.721763

**Authors:** Alma L. Reyes, Amanda H. Rawstern, Elizabeth H. Boughton, Yuxi Guo, Lydia Landau, Jiangxiao Qiu, Michelle E. Afkhami

## Abstract

Global change drivers are reshaping agroecosystems and their sustained functions worldwide. While soil microorganisms underpin the resilience of these systems, the individual and interactive effects of multiple anthropogenic stressors on microbial community structure and function using large-scale field experiments remain poorly understood. Here, we utilize a full-factorial field experiment in a subtropical agroecosystem to investigate how land-use intensity, cattle grazing intensity, and altered precipitation regimes interact to shape soil microbiomes. Combining microbiome sequencing with network analyses and functional bioinformatics, we evaluated effects of these drivers on prokaryotic and fungal diversity, composition, predicted functional profiles, and community structure. Land-use intensity emerged as the primary driver of microbial responses, explaining 25% and 13% of the total variation in community composition for prokaryotes and fungi, respectively. Compared to intensively managed pastures, semi-natural pastures had significantly different community composition for prokaryotes and fungi and exhibited 22% higher fungal diversity. Semi-natural pastures were enriched with decomposer-associated taxa and metabolic pathways related to energy and lipid metabolism indicating enhanced microbial activity. Surprisingly, intensively managed pastures showed higher network modularity but lower network richness, suggesting a trade-off between community compartmentalization and complexity under intensive land management. Grazing and precipitation manipulations induced core microbiome changes within land-use intensities but had no impact on overall community structure and no significant interactions with land-use. Together, these findings suggest that long-term land-use legacies exert a persistent influence on soil microbial community structure, function, and organization, shaping the context within which other global change drivers operate in subtropical agroecosystems.

## 1. Introduction

Global change encompasses a suite of concurrent drivers—including land-use change, climate change, CO_2_ enrichment, nitrogen deposition, and exotic species invasions—that collectively reshape ecosystem processes worldwide (Sage, 2020; Sala et al., 2000; Tylianakis et al., 2008). These global change drivers disrupt ecological functioning by altering species composition (Suleiman et al., 2022), nutrient fluxes (Richardson et al., 2025), and ecosystem stability (Canelas & Pereira, 2022). Land-use change, one of the primary drivers of biodiversity loss (Davison et al., 2021; Newbold et al., 2015), contributes to habitat fragmentation through habitat loss (Haddad et al., 2015) and soil degradation through land conversion (Borrelli et al., 2017). Rising temperatures increase drought severity (Gebrechorkos et al., 2025), shift species’ geographic ranges (Pecl et al., 2017), and disrupt phenological timing (Muluneh, 2021; Parmesan, 2006). Moreover, elevated nitrogen inputs modify plant competitive dynamics (X. Guo et al., 2023), accelerate nutrient imbalances (Peñuelas & Sardans, 2022), and affect biogeochemical cycles (Lu et al., 2011). Collectively, the outcomes of these drivers result in unprecedented stress on natural ecosystems and reduce their capacity to maintain essential ecological functions (e.g., nutrient cycling, primary productivity, community stability), thereby lessening the beneficial ecosystem services they can provide. Ultimately, these pressures act across ecosystems of all types, leading to losses in biodiversity and productivity.

Among these impacted ecosystems, agroecosystems are especially important for feeding human populations, and they are not insulated from the broader global change pressures reshaping natural ecosystems. Approximately 40% of the Earth’s terrestrial land surface is dedicated to agroecosystems (Lal, 2018; Power, 2010), making these human-managed systems a dominant component of the terrestrial biosphere. Agroecosystems supply food, fiber, and livestock resources to sustain the growing global population, generating internal services such as microbial nutrient cycling and soil fertility while also depending on external inputs such as pollination and biological pest control from natural habitats (Power, 2010). While agroecosystems are intentionally managed and experience substantial human driven alterations (e.g., nutrient leaching, soil erosion, water contamination), many global change drivers can exacerbate vulnerabilities created through agricultural management. For instance, agroecosystems can experience intensive tillage, fertilization inputs, and livestock grazing pressures as a component of their land management, which can degrade soil structure and thereby reduce biodiversity (Eldridge & Delgado-Baquerizo, 2017; Hartmann & Six, 2022).

However, the external pressures placed on agroecosystems, such as altered precipitation regimes, warming temperatures, and pests, and other disturbances can compound these problems by pushing managed systems beyond their capacity for resilience. Therefore, even though agroecosystems are anthropogenically altered, global change drivers impose additional stressors that can threaten their productivity, stability, and long-term capacity to support the growing human population.

A major determinant of how agroecosystems respond to such intensifying pressures is the diverse set of belowground processes driven by soil microorganisms, which are highly responsive to environmental shifts and essential to all ecosystems. In agroecosystems, these microbial communities regulate nutrient cycling, store carbon, support plant productivity, and contribute to soil structure, hence making them central to the functioning of these managed systems (Hartmann & Six, 2022; Xiong & Lu, 2022). Soil microorganisms can also suppress pathogens, but under stress, their suppressive ability is diminished thus resulting in pathogen proliferation (Döring et al., 2020; Du et al., 2025). Yet the effects from global change drivers can influence their ability to fulfill these functions under these growing pressures. For instance, altered precipitation patterns can shift the relative abundance of bacteria and fungi in soils, with some studies showing increased precipitation favoring fungal dominance (S. Xu et al., 2020; X. Yang, Zhu, et al., 2021), while land-use change modifies soil properties, such as nutrient availability (Lauber et al., 2008; Leul et al., 2023). Similarly, nutrient enrichment and warming can further influence microbial activity and composition by modifying resource inputs and soil physicochemical conditions (Leff et al., 2015; Metze et al., 2024). Overall, these drivers can reorganize microbial communities and disrupt their capacity to sustain critical agroecosystem functions (Liu et al., 2024).

However, despite the ubiquity of co-occurring global change factors, limited research has investigated how multiple global change factors interact or if factors of a large effect outweigh less stressful factors to influence soil microbial communities (Rillig et al., 2019; Tang et al., 2025; G. Yang et al., 2022). While observational studies have correlated global change factors with natural variation in microbial communities (Bahram et al., 2018; Zhou et al., 2020), and some multiple global change factor experiments in laboratory conditions (Rillig et al., 2019; Rodríguez Del Río et al., 2025), few studies have explicitly investigated these interactions in a manipulative factorial field experiment. Such field experiments are critical for both establishing causality and understanding possible interactive effects of global change on soil microbial communities. Moreover, research assessing microbial responses to environmental change through field experiments has largely focused on temperate systems (Birgander et al., 2018; S. Ma et al., 2018; Matulich et al., 2015), whereas subtropical systems—where the nature and ecological manifestations of climate change are likely to differ substantially—remain understudied. Given that soil microorganisms drive key ecosystem processes (e.g., nutrient mineralization, organic matter decomposition, and carbon sequestration), their comprehensive responses to multiple drivers are central to predicting ecosystem function under intensifying change.

A full understanding of these responses requires not only examining changes in community diversity and composition but also evaluating how relationships among taxa reorganize under these growing pressures. For example, microbial co-occurrence networks provide an additional lens for understanding microbial communities as network properties provide important insight into community structure and dynamics (Byers et al., 2024; Hernandez et al., 2021). These network properties include modularity, which is a measure of how compartmentalized communities are into distinct clusters of taxa (Hernandez et al., 2021; Ling et al., 2016), and the ratio of negative to positive cohesion, which quantify associations among taxa and can provide insight into potential competitive and facilitative interactions among taxa (Hernandez et al., 2021; Herren & McMahon, 2017). Ecological theory suggests that higher modularity and a higher ratio of negative to positive interactions are more stable because if taxa are lost due to disturbance, only their own modules are impacted and not the community at large, therefore fewer positive interactions limit simultaneous species loss (Afkhami et al., 2026; Kajihara & Hynson, 2024). This theory has been experimentally confirmed, showing that microbiomes with high modularity and negative to positive cohesion had increased resistance (i.e., limited compositional shifts after environmental disturbances) and/or recovery ability (i.e., more likely to return to their equilibrium composition after a disturbance)(Agler et al., 2016; Coyte et al., 2015; Herren & McMahon, 2017; Maurice et al., 2024). Therefore, evaluating these network metrics alongside microbial diversity and composition is necessary to fully assess soil microbiomes’ capacity to sustain ecosystem functioning and their resilience to external stressors in extensive and sensitive agroecosystems.

To investigate how soil microbial communities and their network properties respond to multiple global change drivers in agroecosystems, we used a full-factorial field experiment in subtropical humid grasslands of south-central Florida, USA, manipulating land-use intensity, cattle grazing intensity, and precipitation regimes (amount and seasonality). Here, we asked: (1) How do land-use intensity, grazing intensity, and altered precipitation affect soil microbial diversity and composition? (2) How do these factors influence the predicted functional profiles of microbial communities? and (3) Does land-use intensity alter the network structure and stability of microbial communities? By combining high-throughput sequencing of the archaeal and bacterial 16S ribosomal RNA (16S rRNA) gene and the fungal ribosomal internal transcribed spacer (ITS) region, we evaluated how these interacting drivers shape community diversity, composition, functional profiles, and network organization. We hypothesized that land-use intensity, cattle grazing intensity, and precipitation regimes would interact to alter soil microbial community structure, with potential cascading effects on the predicted functional profiles and network organization of communities depending on the magnitude of those structural changes. Moreover, we expected that long-term land-use history would exert the strongest immediate effect through differences in vegetation diversity and nutrient inputs, while cattle grazing intensity and altered precipitation regimes would produce shifts in functional profiles through their influence on soil disturbance and moisture availability. Together, our research aimed to improve our understanding of how interacting global change drivers shape the soil microbiome and influence its capacity to sustain ecosystem functioning in subtropical agroecosystems.

## 2. Materials and Methods

### 2.1. Study area

Our study was conducted at Archbold Biological Station’s Buck Island Ranch, a 4,290-ha commercial cattle ranch part of the United States Department of Agriculture Agricultural Research Service Long-Term Agroecosystem Research Network, located in south-central Florida, USA (27°09′ N, 81°11′ W; Fig. 1). The region experiences a humid subtropical climate with distinct dry (November–May) and wet (June–October) seasons. The mean annual rainfall is 1320 mm, 75% of which primarily falls in the wet season, and the mean annual temperature is 22°C (Boughton et al., 2025). Buck Island Ranch contains two primary land-use intensities that reflect historical differences in land management: intensively managed and semi-natural pastures. Intensively managed pastures are heavily ditched and drained, are dominated by the non-native bahiagrass (*Paspalum notatum*), and support higher cattle stocking rates, particularly during the wet season. In addition, intensively managed pastures were historically fertilized with ∼40 kg P_2_O_5_ ha^-1^ annually from the 1960s until 1986, leaving a legacy of elevated soil phosphorus (Boughton et al., 2025; Swain et al., 2013). Semi-natural pastures, by comparison, are less intensively ditched, have no history of fertilization, support lower cattle stocking rates, and are grazed primarily during the dry season. Additionally, semi-natural pastures are dominated by a mixture of non-native bahiagrass and native grasses such as carpetgrass (*Axonopus* spp.), purple bluestem (*Anatherum cretaceum*), and redtop panicum (*Panicum rigidulum*).

**Figure 1.**
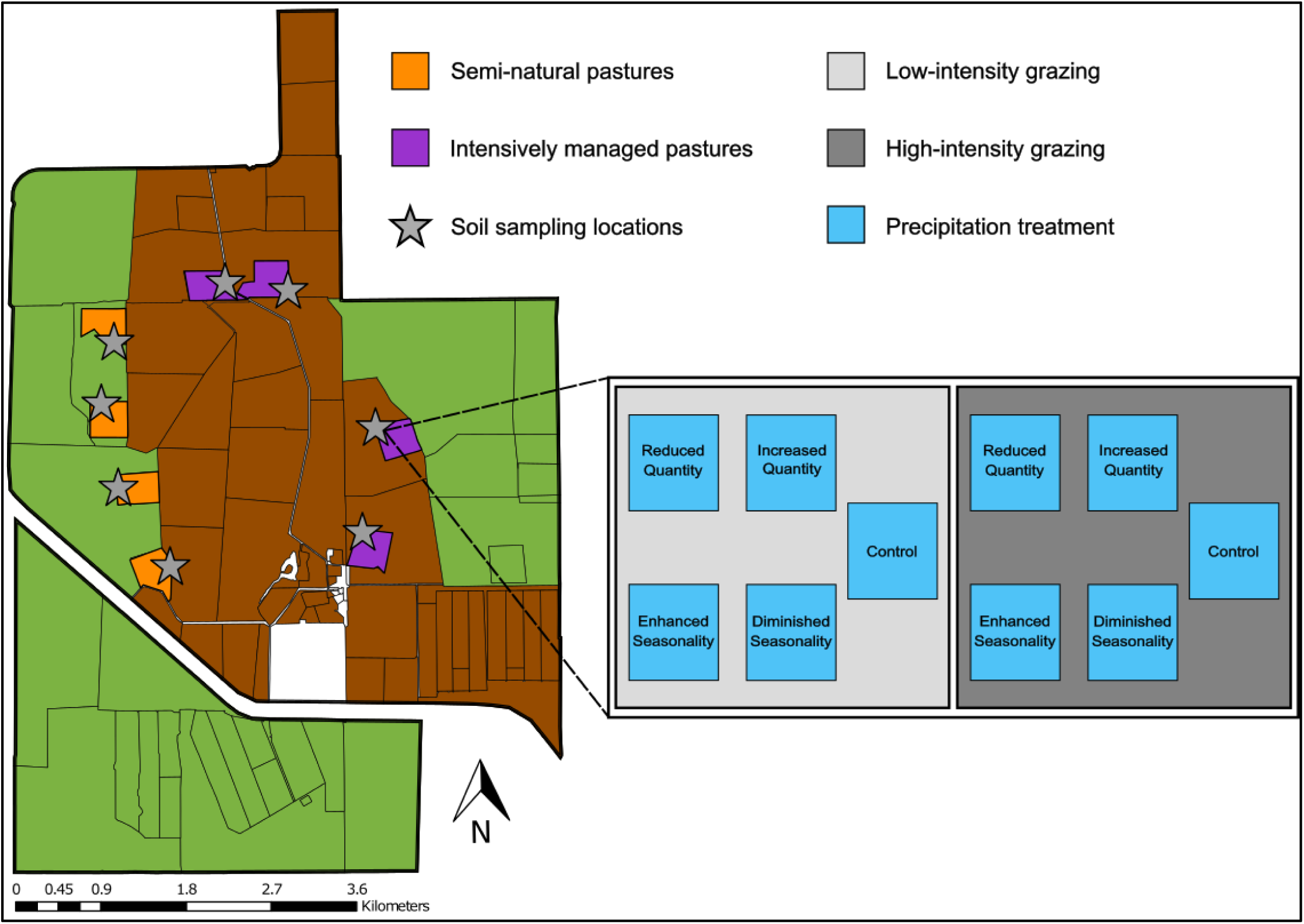
Map of global change factorial experimental at Archbold Biological Station’s Buck Island Ranch. Shaded areas indicate semi-natural pastures (dark orange) and intensively managed pastures (dark orchid). Stars denote the soil sampling locations used in the 2 x 2 x 5 factorial design. Each pasture contains two grazing intensity treatments (low and high), with five precipitation manipulation treatments nested within each grazing level. Note that pastures from each land-use treatment were positioned to match hydrological and topographical conditions.

### 2.2. Experimental design and soil sampling

To investigate the interactive effects of land-use intensity, cattle grazing intensity, and altered precipitation regimes, our research was conducted within an ongoing experiment with a randomized block design (2 x 2 x 5 full-factorial) with four blocks (N = 4) established at Archbold’s Buck Island Ranch. Blocks are positioned along a hydrological and topographical gradient, and each block contains one intensively managed pasture and one semi-natural pasture totaling eight pastures across the experiment. Each pasture is subdivided into two 0.06-ha plots to implement high- and low-intensity cattle grazing treatments. Within each grazing intensity treatment, five plots (4.7-m x 3.1-m) are assigned to one of five precipitation treatments: enhanced seasonality, diminished seasonality, increased quantity, reduced quantity, or an ambient control.

For the grazing intensity treatment, grazing events occur twice per year (April–May and November–December), with livestock density and grazing duration adjusted seasonally for each land-use intensity to maintain target animal use days per hectare (AUD/hectare) that are based on typical AUD/hectare for Buck Island Ranch and the region. Under this framework, distinct stocking rates across land-use intensity and grazing treatments are used to reflect real-world land management practices for this study region. Intensively managed pastures support higher stocking rates annually, ranging from 450–675 AUD/hectare under high intensity and 222–289 AUD/hectare under low intensity while semi-natural pastures support lower stocking rates annually, ranging from 230–289 AUD/hectare under high intensity and 111–124 AUD/hectare under low intensity (Y. Guo et al., 2025). Grazing treatment effects are quantified by measuring vegetation heights immediately before and after each grazing event. Averaged across the grazing events that occurred prior to this study, vegetation heights in intensively managed pastures decreased by 54% under high-intensity grazing and 28% under low-intensity grazing, while in semi-natural pastures, vegetation heights decreased by 35% under high-intensity grazing and 31% under low-intensity grazing (Y. Guo et al., 2025).

Precipitation treatments nested within grazing treatments manipulate rainfall across the wet and dry seasons using rainout shelters and irrigation systems. Treatments are designed to reflect the historical variability and extremes of seasonal precipitation, follow established recommendations for precipitation manipulation experiments, and incorporate downscaled regional climate projections for this region (Beier et al., 2012; Infanti et al., 2020; Knapp et al., 2015). Specifically, the precipitation treatments include: (1) enhanced seasonality, increasing wet season rainfall by 50% while reducing dry season rainfall by 75%; (2) diminished seasonality, reducing wet season rainfall by 50% and increasing dry season rainfall by 75%; (3) increased quantity, involving a 50% increase in wet season rainfall and a 75% increase in the dry season; (4) reduced quantity, comprising a 50% reduction in the wet season and a 75% reduction in the dry season; (5) control, maintains ambient conditions. Regional projections under high-emission scenarios for this region suggest a drying trend, yet high uncertainty remains regarding changes in both total precipitation and its seasonal distribution (Infanti et al., 2020; Stefanova et al., 2012; J. Wang & Kotamarthi, 2015). Seasonality is particularly important in this region, as ecosystem processes (e.g., soil moisture dynamics, plant productivity, and forage quality) can depend on the timing and amount of precipitation, which can disproportionally affect ecological responses compared to changes in total precipitation alone. Therefore, to capture these projections, the diminished seasonality and reduced quantity treatments represent the slightly likely drier futures while the enhanced seasonality and increased quantity treatments represent alternative scenarios characterized by amplified extremes.

In June 2023, one year after experimental manipulations were implemented, we sampled soil microbiomes from all 80 precipitation plots. For each plot, a composite soil sample was collected by homogenizing four soil cores (10-cm depth, 2-cm diameter) collected randomly from points within an ∼58-cm radius from the center of the plot. Samples were transported on ice to the University of Miami and stored at −80°C until DNA extraction.

### 2.3. Soil DNA extraction, amplicon sequencing, and data processing

DNA was extracted from 0.50 g of each soil sample using the DNeasy PowerSoil Pro Kit (Qiagen, 47016). DNA concentrations were quantified using a Qubit 4 Fluorometer. To remove PCR inhibitors, samples were run on a 1.5% agarose gel, and genomic bands were extracted and purified using the E.Z.N.A gel extraction Kit (Omega Bio-Tek, 101318-972). Manufacturer instructions were followed for both extraction and purification protocols. Amplicon libraries were prepared using a two-step dual indexing protocol (Gohl et al., 2016) with magnetic bead cleanups after each PCR step. For amplification, primer pairs 515F/806R and ITS1F/ITS1R were used to target the 16S rRNA gene (archaea/bacteria) and the ITS region (fungi), respectively (Revillini et al., 2022). Evaluation of DNA for quality control were performed using 1% agarose gel electrophoresis. Negative controls with ultrapure water were included to check for contamination. Amplicon libraries were sequenced at the sequencing core at the University of Miami for Genome Technology (Miami, FL, USA) on the Illumina MiSeq platform (v3 PE300).

Sequences were imported and processed using the QIIME2 pipeline (v.2023.9; Bolyen et al., 2019), with optimal trimming parameters determined using FIGARO (Weinstein et al., 2019). The DADA2 algorithm (Callahan et al., 2016) was used to denoise 16S and ITS forward reads (Khan et al., 2022; Lozano et al., 2024; Walsh et al., 2024). Denoised reads were then grouped into amplicon sequence variants (ASVs; 100% sequence similarity) and low-abundance ASVs (<0.25%) were subsequently removed to eliminate spurious sequences (Reitmeier et al., 2021). Representative sequences were aligned with MAFFT (v.7; Katoh & Standley, 2013), and an unrooted phylogenetic tree was constructed using FastTree2 (Price et al., 2010) and midpoint-rooted for phylogeny-based analyses (e.g., weighted UniFrac). Taxonomy was assigned using pretrained naïve Bayes classifiers (Bokulich et al., 2018) trained on the SILVA (prokaryotic; v.138; Quast et al., 2012) and UNITE (fungal; v.10.0; Nilsson et al., 2019) reference databases. ASVs were classified into reference sequence clusters representing approximate species-level groups defined at ≥ 99% sequence similarity.

### 2.4. Data analysis

To assess how land-use intensity, grazing intensity, and precipitation regimes alter the alpha diversity (Shannon diversity index, richness, and evenness) of soil microbial communities, we used generalized linear mixed models implemented in the R package *glmmTMB* (Brooks et al., 2017). Alpha diversity (hereafter “microbial diversity”) indices for ASVs were calculated in the R package *vegan* (v.2.7-1; Oksanen et al., 2025). Land-use intensity, grazing intensity, precipitation treatments, and their interactions were included as fixed effects with block as a random effect. Model residuals were inspected for normality and homoscedasticity using the R package *DHARMa* (v.0.4.7; Hartig, 2024), and the appropriate error distribution family was selected for each response variable. The significance of fixed effects was determined using Type III Wald chi-square tests, followed by Tukey’s HSD post-hoc comparisons using the *emmeans* R package (v.1.11.2-8; Lenth, 2025) when significant effects were detected.

To identify the core prokaryotic and fungal microbiomes across land-use intensity and within grazing treatments, the R package *microbiome* (Lahti & Shetty, 2012) was used. Core taxa were defined as those that maintain a 0.10% relative abundance detection threshold and occur in > 60% of all samples following (Revillini et al., 2023). To evaluate differences in prokaryotic and fungal community composition across treatments, we performed PERMANOVA with 999 permutations with the *adonis2* function (*vegan*; Oksanen et al., 2025), using weighted UniFrac and Bray–Curtis dissimilarity matrices, respectively, with permutations stratified by block. Random forest models with Boruta feature selection (*Boruta*; v.9.0.0; Kursa & Rudnicki, 2010) were subsequently used to identify microbial families important for distinguishing between land-use intensity, grazing intensity, and precipitation treatments.

Predicted functional annotations of prokaryotes were generated using PICRUSt2-SC (v.2.6.2; Wright & Langille, 2025), while fungal trophic modes and ecological guilds were assigned using FUNGuild (v.1.1; Nguyen et al., 2016) based on the species-level taxonomic assignments obtained from UNITE. To assess differences in fungal functional guilds (e.g., wood saprotroph, plant pathogen), the dataset was filtered to retain only taxa with “Highly Probable” and “Probable” confidence rankings, and ASVs not identified at the species level were excluded. From the resulting dataset, we calculated guild richness and identified the three most common guilds using normalized ASV counts. Treatment effects on functional guilds were analyzed using the same generalized linear mixed model structure described above. Predicted prokaryotic functional profiles were generated as Kyoto Encyclopedia of Genes and Genomes (KEGG) Orthologs (KOs), which were subsequently annotated and aggregated into KEGG pathways using the *pathway_annotation* function (*ggpicrust2*; v.2.5.10; C. Yang et al., 2023). To test if prokaryotic functional profiles (KEGG pathways) differed across treatments, a PERMANOVA with 999 permutations on a Bray–Curtis distance matrix was performed, with permutations stratified by block. Differential abundance analyses of microbial taxa and prokaryotic functional KEGG pathways were performed using analysis of compositions of microbiomes with bias correction 2 (ANCOM-BC2; Lin & Peddada, 2024) implemented in the R package *ANCOMBC* (v.2.4.0; Lin et al., 2022; Lin & Peddada, 2020) with false discovery rate correction. Features (i.e., taxa and KEGG pathways) were excluded by prevalence, with a 10% cutoff for prokaryotes and KEGG pathways and a 20% cutoff for fungi.

To examine if and how land-use and precipitation within each land-use intensity affected microbial community structure, co-occurrence networks were constructed separately for prokaryotic and fungal communities using FastSpar (v.1.0.0; Watts et al., 2019), a parallel implementation of SparCC (Friedman & Alm, 2012). Samples were aggregated by land-use intensity and precipitation treatment, resulting in one network per combination for each microbial group (prokaryotes and fungi), totaling 10 networks per group. Within each treatment combination, ASVs were filtered to retain taxa present in ≥ 5% of samples and occurring in more than two replicates; peripheral taxa (e.g., nodes with low connectivity) were excluded (Rawstern et al., 2025). Pairwise correlations were inferred with 1,000 bootstrapped iterations, and only significant associations (*p* < 0.05) were retained. For each network, the structural properties modularity (i.e., community compartmentalization), negative:positive cohesion ratio (i.e., associations among taxa), and network richness (i.e., number of nodes with moderately- and highly-connected taxa) were quantified using the NetworkX python package (v.3.4.2; Hagberg et al., 2008). A MANOVA was used to test the effects of land-use intensity and microbial group on network structure with network metrics treated as a multivariate response and follow-up ANOVAs of individual metrics were performed. Statistical analyses were performed in R v.4.3.2 (R Core Team, 2023).

## 3. Results

### 3.1. Less intensive land management increases fungal, but not prokaryotic, microbial diversity

Fungal diversity, richness, and evenness were more responsive to the experimental treatments than prokaryotes. In particular, we found that fungal diversity was ∼22% greater in semi-natural pastures compared to intensively managed pastures (χ²(1, N = 4) = 9.00, *p* = 0.003; Fig. 2A,B). The difference was primarily driven by a ∼26% increase in fungal richness (χ²(1, N = 4) = 5.09, *p* = 0.024; Fig. 2C,D) and ∼13% increase in fungal evenness in semi-natural pastures (χ²(1, N = 4) = 9.91, *p* = 0.002). By contrast, prokaryotic diversity and richness were not affected by any of the treatments. For both fungi and prokaryotes, there were no significant interactive effects of land-use intensity, grazing intensity, and precipitation treatments on these alpha diversity metrics (Table S1).

**Figure 2.**
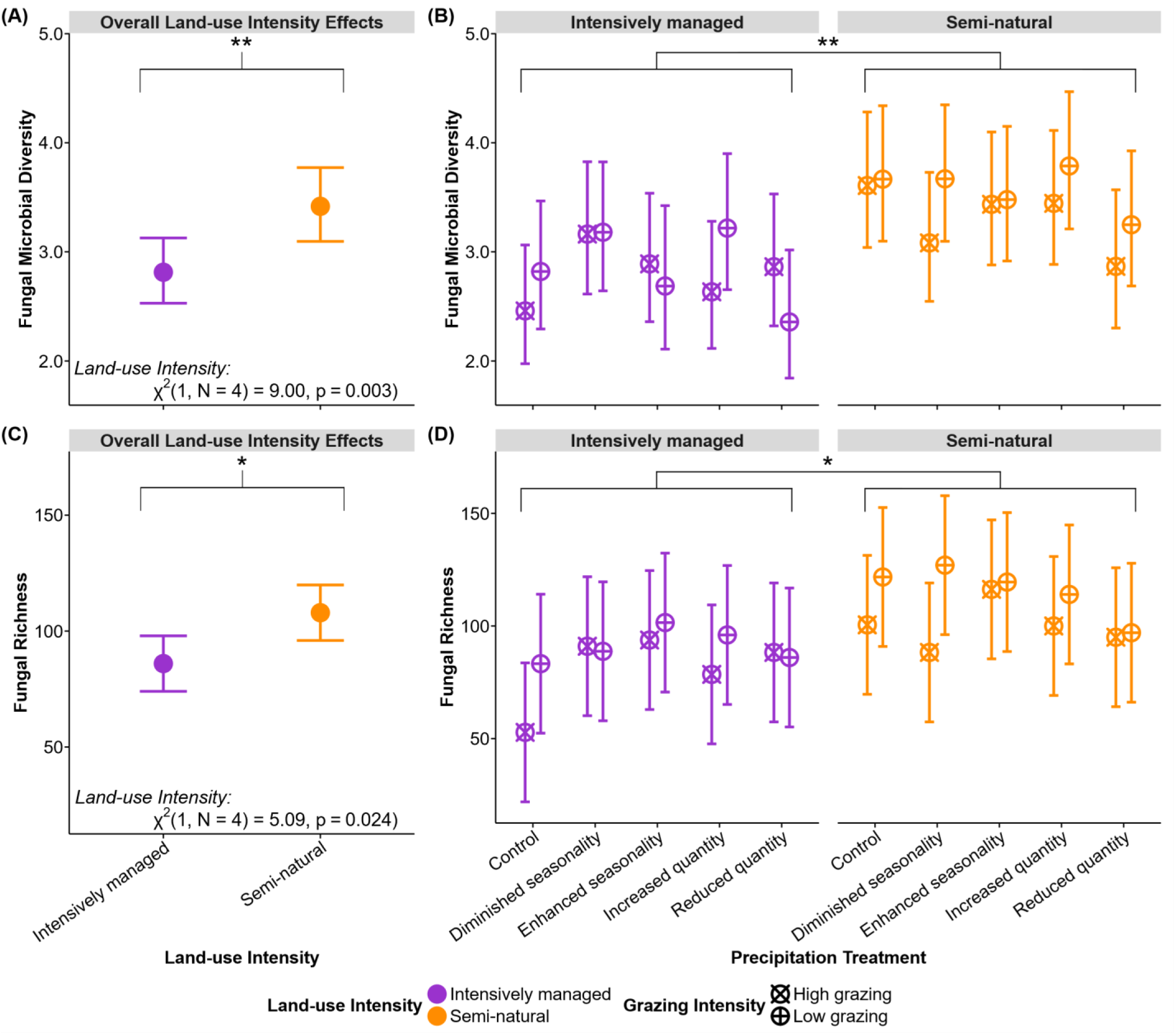
Land-use intensity affects fungal microbial diversity with semi-natural pastures exhibiting higher fungal diversity than intensively managed pastures. (A, C) Overall fungal microbial diversity (Shannon diversity index) and richness across land-use intensity. (B, D) Fungal diversity and richness across land-use intensity, grazing intensity, and precipitation treatments. Values represent mean ± SE. Pairwise differences were determined by Tukey-adjusted post hoc tests. * indicates *p* ≤ 0.05; ** indicates *p* ≤ 0.01.

### 3.2. Land-use intensity significantly altered microbial community composition

Across all treatments examined, the strongest driver of differences in prokaryotic and fungal taxonomic community composition was land-use intensity (prokaryotic: pseudo-*F* = 24.93, *p* < 0.001; fungi: pseudo-*F* = 11.29, *p* < 0.001; Fig. 3A,B, S1, S2), with no significant main or interactive effects of grazing intensity and precipitation treatments on overall community composition (Table S2).

**Figure 3.**
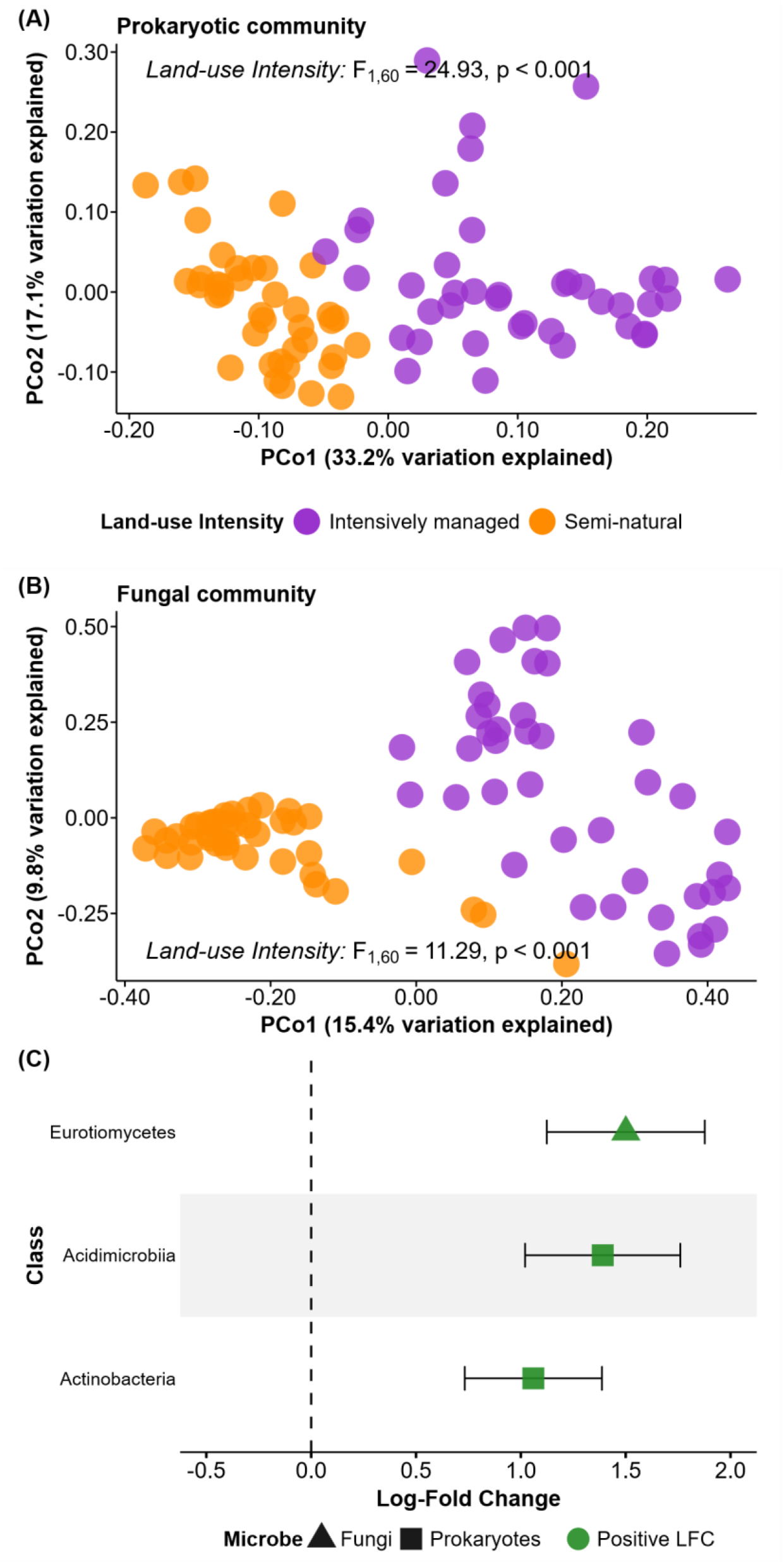
Land-use intensity shapes prokaryotic and fungal community composition in intensively managed and semi-natural pastures. Principal coordinate analysis of (A) prokaryotic community composition (weighted UniFrac) (B) fungal community composition (Bray–Curtis dissimilarity) and (C) Differentially abundant microbial classes identified by ANCOM-BC2 in the land-use intensity comparison. Intensively managed pasture was used as the reference level. Positive values indicate higher abundance in semi-natural pastures. Symbols indicate microbial group and colors denote positive or negative log-fold change (LFC) relative to the reference (intensively managed pastures). Symbols represent mean LFC and error bars represent SE from ANCOM-BC2.

To further characterize key changes in community composition, random forest models identified eight prokaryotic and 30 fungal families as important for distinguishing between land-use intensity (Table S3). Notably, the majority of these taxa (six prokaryotic and 23 fungal) were more abundant in semi-natural, with the remainder associated with intensively managed land-use. In particular, the family Herpotrichiellaceae (which includes saprotrophic, pathogenic, and endophytic fungi) was among those enriched in semi-natural pastures. For grazing intensity, two fungal families were identified as important for distinguishing between low- and high-grazing treatments (Table S4). Specifically, the saprotrophic family Didymosphaeriaceae and an unclassified family in the phylum Basidiomycota were more abundant in low grazing treatments. However, no prokaryotic families were identified as important for distinguishing between grazing intensity treatments. Precipitation treatments revealed a parallel pattern with two distinct taxa (Table S5), where the prokaryotic family Bacillaceae (nutrient cyclers involved in plant development) was more abundant in the diminished seasonality treatment, while an unidentified fungal Ascomycota family showed the highest abundance in the control treatment.

Differential abundance analysis revealed 3 microbial classes that significantly changed in relative abundance in response to land-use intensity (2 prokaryotic, 1 fungi; Fig. 3C). Among prokaryotes, the decomposers Acidimicrobiia and Actinobacteria showed the most pronounced abundance increase in semi-natural pastures (approximately +1–1.4 log-fold change (LFC); ∼2.7–4x higher abundance), which was mirrored by a 1.5 LFC (∼4.5x) increase in a fungal decomposer (class Eurotiomycetes) in semi-natural pastures. Conversely, no significant differences in the relative abundance of prokaryotic and fungal classes were detected for the grazing intensity and precipitation treatments.

### 3.3. Grazing intensity and precipitation shift core microbiomes within land-use intensity

While land-use intensity was the dominant driver of microbial diversity and community composition, grazing intensity and altered precipitation regimes shifted core microbiome membership within each land-use intensity (Fig. S3-S10). Within intensively managed pastures, several important prokaryotic families varied in core membership across treatments. For example, Methanobacteriaceae, hydrogenotrophic methanogens, were core members under low-grazing intensity across all precipitation treatments (Fig. S5). However, under high grazing, Methanobacteriaceae was absent in the reduced quantity precipitation treatment, suggesting intense grazing and water limitation constraints methanogenic niches (Fig. S6). In addition, Chitinophagaceae, a family of chitin- and cellulose-degrading bacteria associated with plant residues and soil fertility, were identified as core members exclusively in intensively managed pastures. In contrast, Acetobacteraceae, which are plant-growth promoting and nitrogen fixing bacteria, were a core family unique to semi-natural pastures. Despite differences in core prokaryotic families, several prokaryotic families were shared across land-use intensities (Fig. 4A,B). Notably, Xanthobacteraceae, a metabolically versatile group capable of degrading complex organic compounds and aiding in plant development, was identified as part of the core microbiome across treatments in both land-use intensities.

**Figure 4.**
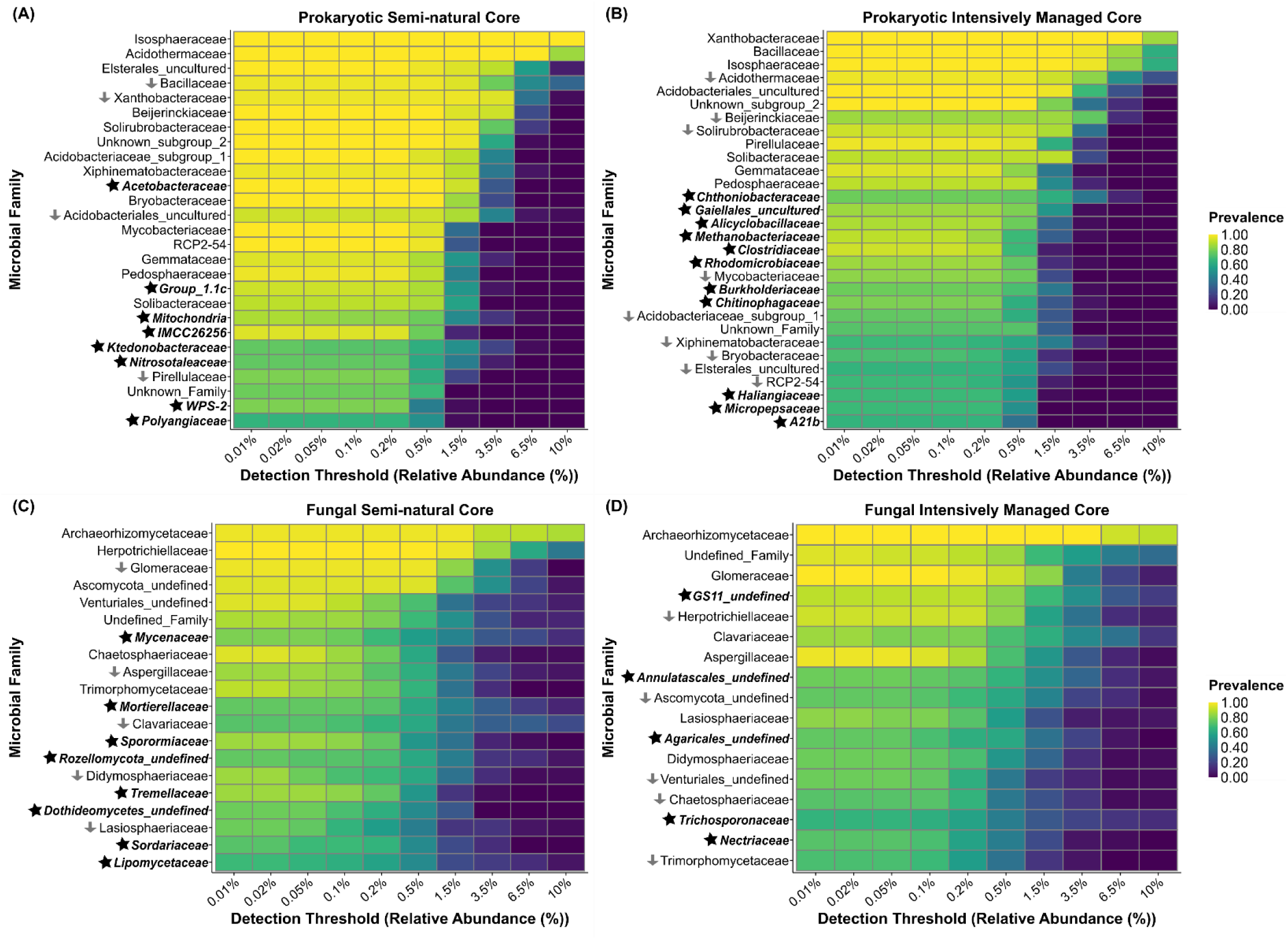
Land-use intensity shifts core membership of microbial families. Core families are organized by relative abundance detection thresholds and shaded by prevalence, with upper and lower panels representing prokaryotic and fungal families, respectively. Stars represent families unique to each land-use intensity, and arrows indicate families that decreased in prevalence.

Fungal communities showed similar treatment-dependent changes in core membership. In semi-natural pastures, Clavariaceae, a group of root-associated saprotrophs, was identified as a core family under low-intensity grazing (Fig. S7). By contrast, under high-grazing intensity, Clavariaceae fell below detection and prevalence thresholds in the reduced quantity and diminished seasonality treatments, indicating sensitivity to combined grazing and precipitation changes (Fig. S8). Several fungal families were consistently observed across land-use intensities, including Glomeraceae (arbuscular mycorrhizal fungi) and Archaeorhizomycetaceae (rhizosphere-associated)(Fig. 4C,D). In addition to these shared fungal families, Mortierellaceae, a family of copiotrophs capable of degrading hemicellulose and chitin and contributing to carbon cycling, was identified as a core family present only in semi-natural pastures.

A full list of the core prokaryotic and fungal families within each treatment combination available in Fig. S3–S10.

### 3.4. Functional predictions of prokaryotes and fungi show distinct responses to land-use, grazing-intensity, and precipitation

Predicted functional profiles of prokaryotic communities were influenced exclusively by land-use intensity (pseudo-*F* = 23.35, *p* < 0.001; Fig. 5; Table S6). Differential abundance analysis identified 79 KEGG pathway classes that responded significantly to land-use intensity (Fig. S11). Of these, 40 exhibited positive LFCs, indicating higher relative abundance in semi-natural pastures. These pathways were primarily associated with metabolic categories – e.g., biosynthesis of secondary metabolites, lipid metabolism, energy metabolism – with effect sizes around +0.24 LFC (∼1.3x higher abundance in semi-natural pastures; Fig. S11). The remaining 39 pathway classes showed negative LFCs near −0.31 (∼1.4x more abundant in intensively managed pastures: Fig. S11) and largely corresponded to organismal systems (e.g., excretory and digestive system).

**Figure 5.**
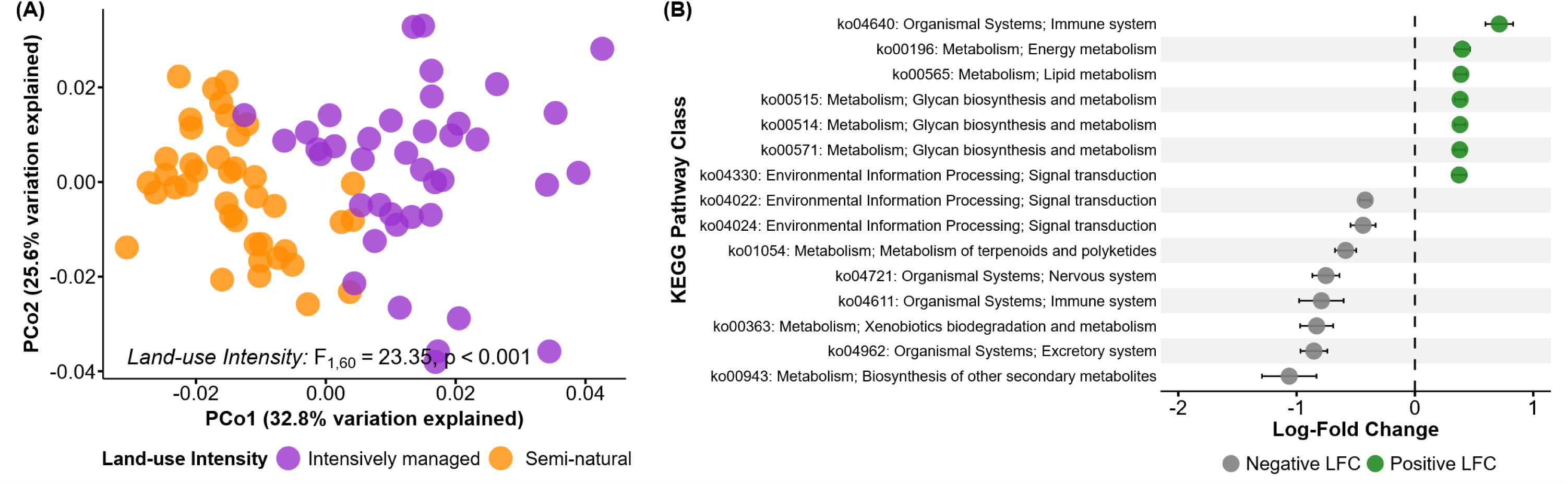
Land-use intensity affects prokaryotic functional KEGG pathways in intensively managed and semi-natural pastures. (A) Principal coordinate analysis of KEGG pathway abundances based on Bray–Curtis dissimilarities. (B) Differentially abundant KEGG pathway classes identified by ANCOM-BC2 in the land-use intensity comparison (15 most significant pathways shown). Intensively managed pasture was used as the reference level. Positive values indicate higher abundance in semi-natural pastures. Colors indicate whether KEGG pathway classes were enriched or depleted relative to the reference. Points represent mean log-fold changes, and error bars represent SE from ANCOM-BC2.

Functional guilds of the fungal communities in our experiment, while most frequently influenced by land-use changes, were also significantly affected by interactions among multiple global change factors. We identified 23 functional guilds in our samples, with “undefined saprotroph” (59.7%), wood saprotroph (6.1%), plant saprotroph (5.6%) most common based on relative abundance (Fig. S12). Overall guild richness varied with land-use intensity; semi-natural pasture had ∼18% higher richness than intensively managed pastures (χ²(1, N = 4) = 4.23, p = 0.040). While we did not detect significant effects of the other treatments or their interactions on guild richness, multiple factors emerged as important for changes in the abundances of the three most common guilds (Table S7). Similar to guild richness, the group of “undefined” saprotrophs (i.e., saprotrophs without detailed information on food source preference) were affected solely by land-use intensity, showing a ∼31% higher abundance in intensively managed pastures compared to semi-natural pastures (χ²(1, N = 4) = 3.93, p = 0.047). Unlike that broad category of “undefined” saprotrophs, wood saprotroph (i.e., wood-decay fungi) abundance was influenced by the interaction between land-use intensity and precipitation treatment (χ²(4, N = 4) = 12.75, p = 0.013). Specifically, in control precipitation plots, wood saprotrophs were 226% more abundant in intensively managed pastures than semi-natural pastures (p = 0.040), suggesting that under ambient precipitation decomposers of complex carbon sources are enhanced in managed conditions. However, in semi-natural pastures, diminished seasonality significantly increased wood saprotrophs abundance, showing a 483% increase over the reduced quantity precipitation treatment (p = 0.023). Similarly, plant saprotroph (i.e., organisms that break down dead plant material) abundance was affected by land-use intensity (χ²(1, N = 4) = 6.66, p = 0.009) and the interaction between land-use intensity and precipitation treatment (χ²(4, N = 4) = 9.96, p = 0.041). Notably, Tukey-adjusted post-hoc comparisons detected no significant differences for plant saprotroph abundance across the two land-use intensities (p = 0.220), but instead the effect of land-use intensity was contingent on precipitation levels. Specifically, in the control precipitation treatment, semi-natural pastures showed an 188% increase in plant saprotrophs abundance compared to intensively managed pastures (p = 0.016). Furthermore, within intensively managed pastures, enhanced seasonality significantly increased plant saprotrophs abundance by ∼227% compared to the control (p = 0.049).

### 3.5. Microbial community networks differed by land-use intensity

Network structure differed significantly by land-use intensity (*F*_3,14_ = 27.38, *p* < 0.001) and microbial group (i.e., prokaryotic and fungi; *F*_3,14_ = 39.01, *p* < 0.001; Fig. 6; Table S8).

**Figure 6.**
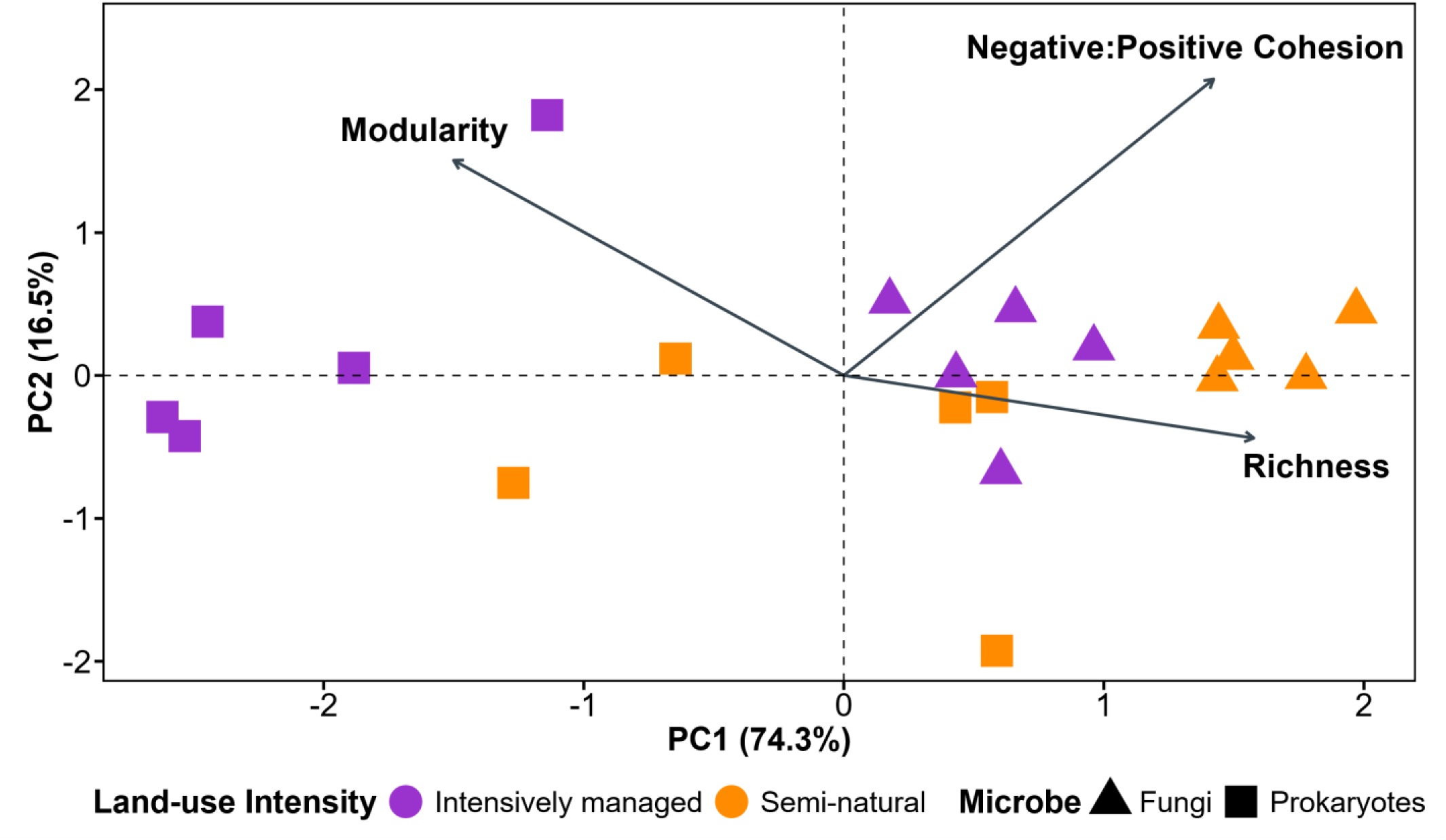
Network properties for prokaryotic and fungal networks differ between land-use intensity. Each point represents a prokaryotic or fungal co-occurrence network. Networks are ordinated by principal components of modularity, negative:positive cohesion, and richness.

Land-use intensity significantly influenced network modularity (*F*_1,16_ = 11.40, *p* = 0.004) and richness (*F*_1,16_ = 65.39, *p* < 0.001), though it had no effect on negative:positive cohesion ratio (*F*_1,16_ = 2.05, *p* = 0.171). Intensively managed pastures had 21% higher network modularity than semi-natural pastures, indicating greater compartmentalization of soil communities in intensively managed land. In contrast, semi-natural pastures had 61% greater network richness compared to intensively managed pastures, emphasizing the greater diversity of microbiomes in semi-natural pastures. When comparing between microbial groups, we found that prokaryotic and fungal networks exhibited distinct structural characteristics across all network metrics. Prokaryotic networks had 21% greater modularity than fungal networks (*F*_1,16_ = 13.07, *p* = 0.002). Fungal networks, by comparison, were characterized by a significantly higher negative:positive cohesion ratio and richness, with a 51% higher negative:positive cohesion ratio (*F*_1,16_ = 19.14, *p* < 0.001) and ∼70% greater richness (*F*_1,16_ = 79.98, *p* < 0.001) relative to prokaryotic networks.

Interestingly, we detected a significant land-use and microbial group interaction for modularity (*F*_1,16_ = 4.76, *p* = 0.044). Specifically, intensive land management had a 33% increase in modularity for prokaryotic networks, while fungal networks had a much weaker increase of 8%, indicating that compartmentalization increases under intensive land management detected in the overall microbial community was largely driven by prokaryotes.

## 4. Discussion

In this study, we examined the individual and interactive effects of land-use intensity, grazing intensity, and altered precipitation on prokaryotic and fungal communities in a subtropical agroecosystem. Across different levels of microbial organization, including taxonomic diversity, community composition, predicted functional profiles, and co-occurrence networks, we found that land-use intensity consistently emerged as the dominant driver of microbial responses. Fungal communities showed sensitivity to land-use intensity through changes across diversity metrics, functional guilds, and co-occurrence networks. By comparison, prokaryotic responses were generally more subtle, characterized by structural and functional changes rather than significant changes in taxonomic diversity.

### 4.1. Land-use intensity as the primary driver of microbial composition

Land-use intensity emerged as the primary factor structuring prokaryotic community composition, though these changes were not mirrored by shifts in microbial diversity (Fig. 3A; Table S1). These findings indicate that land-use intensity drives species turnover without net richness loss, a shift likely mediated by differences in soil physicochemical conditions and vegetation structure (Z. Guo et al., 2022; A. Xu et al., 2021). Our study reveals that land-use intensity alone can drive species turnover in prokaryotic communities, even under experimentally varied grazing intensity and altered precipitation regimes. Previous large-scale field experiments in this subtropical agroecosystem showed that bacterial community composition responded to land-use intensity as well as to the interaction between grazing and fire (Y. Guo et al., 2023), with grazing and fire manipulated as presence-absence treatments rather than intensity gradients. Our work highlights that, even when the overall pattern of land-use intensity driving turnover is similar, manipulating drivers along intensity gradients, rather than as binary presence-absence factors, is important for capturing how microbial communities respond to the degree of land-use change.

By contrast, fungal communities were affected by land-use intensity at both the compositional and diversity level (Fig. 3B; Table S1). Semi-natural pastures supported higher fungal diversity driven by increases in both richness and evenness (Fig. 2), indicating these habitats maintain a larger, more equitably distributed pool of fungal taxa. These differences in fungal microbial communities between land-use intensities are in line with previous work in temperate systems, which observed greater sensitivity of fungal communities to increased land-use intensification (Navarro-Noya et al., 2021; Xue et al., 2023), though contrast with work in tropical systems which showed only compositional changes without diversity loss (e.g., tropical rainforest conversion; Brinkmann et al., 2019; Sun et al., 2020). The contrasting responses between prokaryotes and fungi likely reflect fundamental differences in how these groups respond to disturbance, with fungal communities being more sensitive to land-use driven changes in soil structure and plant cover than prokaryotes. In addition, variable subtropical rainfall potentially favors prokaryotes, whose rapid response to wet-dry pulses can buffer community structure, while fungi dependent on stable hyphal networks are more vulnerable to disruption, delaying detectable directional changes in prokaryotic communities. For instance, prokaryotes possess broad metabolic capabilities and can respond rapidly to shifting environmental conditions (Chen et al., 2021; Wani et al., 2022), potentially allowing an early adjustment to land management that dampens later diversity differences across land-use intensities. Conversely, fungi can exhibit greater sensitivity to intensive land use because they depend on a stable soil matrix. Physical disturbances (e.g., grazing by livestock) can fragment fungal hyphal networks, thereby reducing their biomass and thus affecting efficient nutrient-uptake (Hartmann & Six, 2022; Schimel et al., 2007). Consequently, intensive land management practices that disrupt soil structure and reduce plant diversity can lead to constraints in hyphal connectivity and niche heterogeneity, thus resulting in declines in fungal diversity (Shen et al., 2021; Xue et al., 2023).

Further, the abundance of the decomposer classes Acidimicrobiia, Actinobacteria, and Eurotiomycetes highlights the capacity of semi-natural pastures to support decomposition and nutrient cycling processes. As members of the phylum Actinobacteria, Acidimicrobiia and Actinobacteria contribute to the breakdown of complex organic matter and drive the cycling of nutrients such as carbon, nitrogen, and potassium (Araujo et al., 2020; B. Zhang et al., 2019). Higher plant diversity in semi-natural pastures likely promotes the abundance of these classes by providing more heterogenous litter and organic matter inputs. Similarly, the fungal class Eurotiomycetes are established copiotrophic decomposers known for their ability to process plant-derived organic residues and the recycling of carbon (D. Wang et al., 2025; X. Yang, Liang, et al., 2021). Enrichment of these prokaryotic and fungal decomposer lineages indicate that semi-natural pastures are essential for organic matter turnover. Thus, these less intensively managed pastures (i.e., semi-natural) potentially serve as functional reservoirs within the wider agroecosystem, indirectly increasing the productivity of intensively managed pastures by contributing to balanced nutrient dynamics and ecosystem stability at the landscape scale.

Moreover, grazing intensity and altered precipitation treatments had minimal effects on microbial diversity and composition, at least within the relatively short period over which these two treatments were imposed, and no interactive effects were observed. Our findings suggest that the soil microbiome in this subtropical agroecosystem is more strongly constrained by long-term management legacies rather than by short-term variation in disturbance or changes in rainfall imposed by grazing and precipitation treatments, which may take a longer time to manifest.

Similar resistance to experimental manipulation has been observed in systems with strong historical land management modifications, where microbial communities are more strongly influenced by land use (De Vries et al., 2012; Osburn et al., 2021). Importantly, it should be noted that while grazing intensity and precipitation manipulation were experimentally imposed, land-use intensity reflects long-term management legacies that integrate historical soil disturbance, vegetation composition, and nutrient inputs, which potentially constrain the microbial response to these shorter-term treatments.

### 4.2. Land-use intensity structures the core microbiome, with grazing and precipitation modulating within-land-use-variation

Land-use intensity drove distinct shifts in core microbiome membership (Fig. 4), with intensively managed pastures favoring methanogens and semi-natural pastures supporting nitrogen-fixers and root-associated fungi, with additional variation arising from grazing and precipitation within land-use intensities (Fig. S3-S10). Under intensive land management, prokaryotic core families included Methanobacteriaceae, versatile methanogens that utilize methylotrophic and hydrogenotrophic pathways, which allows them to remain active across a wide range of substrates (Y. Zhang et al., 2019), and Chitinophagaceae, which degrade complex plant polymers (e.g., galactan and arabinan; Huang et al., 2023). Interestingly, the absence of Methanobacteriaceae under combined high grazing and the reduced quantity precipitation treatment suggests that these factors disrupt methanogenic niches even in intensively managed pastures. In addition, Xanthobacteraceae – which can aid in the recycling of nutrients such as carbon and nitrogen (Xie et al., 2023; Y. Yang et al., 2024) – occurred under both land-use intensities but became a dominant member of the core microbiome in intensively managed pastures. The emergence of these prokaryotic families in the intensively managed core suggest selection for taxa adapted to exploiting the high nutrient and fluctuating redox conditions of human-supplemented systems, consistent with observations in similar systems (Chroňáková et al., 2015; Wei et al., 2022).

Fungal core microbiomes revealed similar restructuring in core microbiome complexity under intensive land management. Glomeraceae (arbuscular mycorrhizal fungi; Y. Ma et al., 2021) and Archaeorhizomycetaceae (rhizosphere fungi; Pinto-Figueroa et al., 2019) formed stable fungal core members across land-use intensities, suggesting selective retention of root associated fungi. Additionally, Clavariaceae was a core family across both land-use intensities, though showed sensitivity to high grazing in semi-natural pastures under the diminished seasonality treatment. The changes in fungal core membership highlights how intensive management can narrow the range of fungal families consistently supported across these systems.

Despite land-use intensity primarily structuring prokaryotic and fungal core microbiome membership, grazing and altered precipitation regimes exerted secondary effects that modulated family presence within each land-use intensity—even when these treatments showed no overall effects on community composition. Our work suggests that, although land-use intensity remains the dominant filter, additional environmental drivers can generate detectable signals at finer scales, suggesting gradual community changes over longer timescales.

### 4.3. Functional implications of land-use driven microbial community shifts

Changes in microbial community composition associated with land-use intensity were accompanied by corresponding changes in predicted functional profiles of prokaryotic and fungal microbial communities (Fig. 5, S11, S12). These changes suggest that land-use intensity influences not only taxonomic community structure but also translate into functional differences, indicating that functional overlap among communities was limited across land-use intensities. In prokaryotic communities, functional profiles differed exclusively by land-use intensity, with semi-natural pastures enriched in pathways associated with core metabolic processes, including energy metabolism, lipid metabolism, and secondary metabolite biosynthesis (Fig. 5, S11, Table S6). These functions are commonly associated with metabolically versatile communities capable of processing diverse substrates, as has been observed in less intensively managed systems characterized by greater organic matter heterogeneity and lower nutrient inputs (Delgado-Baquerizo et al., 2020; Leff et al., 2015). Intensively managed pastures displayed distinct patterns, characterized by enrichment in pathways associated with organismal systems (e.g., digestive system, excretory system). These shifts potentially reflect the metabolic processing of manure deposition and other livestock-derived inputs, which has been shown to be a characteristic of systems with high cattle stocking density (Zhao et al., 2022).

Fungal functional guild patterns further reinforce the central role of land-use intensity. Semi-natural pastures supported higher fungal guild richness (Fig. S12; Table S7), consistent with the increased taxonomic diversity which could be a result of higher plant diversity in semi-natural pastures (Boughton et al., 2025). The dominance of saprotrophic guilds across all treatments suggests the importance of fungi as primary decomposers in this system. While differences in relative abundance of wood and plant saprotrophs indicate that land-use intensity alters functional guilds related to decomposition rather than eliminating such roles. Moreover, interactions between land-use intensity and precipitation for specific saprotrophic guilds suggest that fungal functional responses are context-dependent. Management history appears to set the conditions under which altered precipitation regimes influence fungal activity. The potential mechanisms for this could be a result of interactions with plant inputs and litter quality. These findings highlight that functional responses to climate-related drivers may be constrained or amplified by long-term land-use legacies.

The alignment between taxonomic and functional differentiation in this system suggests that functional redundancy among microbial taxa is limited under contrasting land-use intensities, likely mediated by environmental filtering that favors distinct metabolic strategies, further reducing overlap among communities and amplifying functional differentiation. Although functions were inferred from taxonomy using established methods (PICRUSt2-SC and FUNGuild) and thus reflects functional potential rather than measured activity, these tools still provide important insights into shifts in prokaryotic metabolic pathways and fungal functional roles in systems facing complex and multifactor global changes and the multifactorial experiments required to evaluate human-impacts on these systems.

### 4.4. Microbial network reorganization under contrasting land-use intensities

Beyond changes in diversity and function, land-use intensity strongly reorganized microbial co-occurrence networks and these responses differed in prokaryotic and fungal networks (Fig. 6). Unexpectedly, intensively managed pastures had higher network modularity than semi-natural pastures, indicating that microbial taxa in these systems are more strongly compartmentalized into subsets of organisms that tend to co-occur under similar environmental conditions. Our result contrasts with the common expectation that environmental stress reduces network modularity (Felipe-Lucia et al., 2020; Hernandez et al., 2021; Romdhane et al., 2022; Wu et al., 2023) but aligns with observations in other managed system where disturbance increases modularity and network stability (Cornell et al., 2023). Potential mechanisms for this pattern include the long-term fertilization and reduced plant diversity in intensively managed pastures, which can create more uniform soil physicochemical conditions and strongly favor microbial assemblages composed of taxa sharing similar ecological tolerances or metabolic requirements (Schmidt et al., 2019; Q. Xu et al., 2020). In particular, prokaryotic networks in our study had higher network modularity than fungal networks, suggesting that fungal community stability, in addition to diversity and richness, is sensitive to land-use intensity.

In contrast, network richness followed the more conventional stress gradient. Specifically, network richness responded strongly to land-use intensity and differed by microbial groups. Across both prokaryotes and fungi, semi-natural pastures supported higher network richness than intensively managed pastures. For fungal networks, higher network richness observed in semi-natural pastures suggests that diverse plant communities allow more fungal taxa to co-occur. By contrast, prokaryotic network richness was lower than fungal networks overall, but the difference between fungal and prokaryotic richness was smaller in semi-natural than under intensive land management. This pattern indicates a broader range of prokaryotic taxa are supported across heterogenous environments in semi-natural pastures. The reduced richness in intensively managed pastures suggests that long-term management limits the number of prokaryotic taxa that consistently co-occur, potentially through changes in soil conditions. The negative to positive cohesion ratio differed between microbial groups but was not influenced by land-use intensity. For fungal networks, the higher negative to positive cohesion ratio indicates that microbial networks are characterized by more negative associations rather than positive associations, independent of land-use intensity. In contrast, prokaryotic networks exhibited a lower negative:positive cohesion ratio, indicating a greater prevalence of positive association among prokaryotic taxa.

Together, our work reveals a microbial group specific response in subtropical agroecosystems, where prokaryotes gain modular robustness under intensive management, while fungal networks remain susceptible to environmental stress. These contrasting responses shed light into stress-modularity relationships and suggest that microbial organization, and its links to ecosystem stability, varies not just by management but by microbial group and biogeography context.

## 5. Conclusions

The work from this study demonstrates that land-use intensity exerts a dominant and persistent influence on soil microbial communities in subtropical agroecosystems, shaping taxonomic composition, predicted functional profiles, and network organization more strongly than short-term grazing or precipitation manipulations. Fungal communities were particularly sensitive, exhibiting both higher diversity and functional guild richness under lower-intensity land use, while prokaryotic responses to land-use intensity were primarily reflected in functional profiles and network structure differences. The findings from our work suggest that long-term management legacies set the baseline context within which other global change drivers operate, influencing microbial-mediated ecosystem processes within subtropical agroecosystems. Within this context, grazing intensity and altered precipitation regimes modified core microbiome membership, but these changes did not propagate to broader community-level changes.

Through the integration of multifactorial field experiments and analyses of diversity, composition, functional capacity, and microbial networks, our results highlight the importance of considering multiple aspects of microbial organization when predicting agroecosystem resilience and the capacity of soil communities to maintain essential ecosystem function under ongoing global change. Based on these findings, we advocate for future work integrating direct functional assays and multi-omics (e.g., metagenomics, metatranscriptomics) across multiple timepoints.

With the advancement of sequencing technologies making these approaches increasingly possible, these tools can be used to help clarify the mechanistic links between microbial community structure, function, and ecosystem processes in agroecosystems. Overall, our research and future work building on these findings will provide valuable insight into how soil microbial communities respond to multiple global change drivers in a rapidly changing world, which is crucial for effectively managing agroecosystems in the Anthropocene.

## Supporting information

Supplement_TablesS1-S8_and_FiguresS1-S12

## Acknowledgements

The authors would like to acknowledge members of Archbold’s Buck Island Ranch Agroecology Lab for their work with maintaining the rainout infrastructure and Buck Island Ranch Operations for their assistance with cattle movements. We thank Damian J. Hernandez for their guidance during the early stages of this project and the members of the Afkhami and Searcy labs for providing feedback on manuscript preparation. This work was supported by the USDA National Institute of Food and Agriculture [grant numbers 2023-67019-39728; 2021-67013-33617; FLA-FTL-006277]. This research was supported in part by the United States Department of Agriculture Agricultural Research Service (USDA-ARS) Long-Term Agroecosystem Research (LTAR) network. LTAR is supported by the United States Department of Agriculture. Any opinions, findings, conclusion, or recommendations expressed in this publication are those of the author(s) and do not necessarily reflect the view of the U.S. Department of Agriculture. ALR was supported by the Lisa D. Anness Graduate Fellowship and MEA was supported by NSF DEB Awards 2505581, 2030060, and 1922521.

