## Supplement_TablesS1-S8_and_FiguresS1-S12 for "Land-use intensity overrides grazing and precipitation effects on soil microbial communities in a subtropical agroecosystem"

**Table S1.** Results of generalized linear mixed models of land-use intensity, grazing intensity, and precipitation treatment effects on prokaryotic alpha diversity metrics (Shannon diversity, richness, and evenness). Bold values indicate significant effects ($p < 0.05$).

| | | Shannon Diversity | | | Richness | | | Evenness | | |
|---|---|---|---|---|---|---|---|---|---|---|
| | Treatment | Chisquare | df | *p* | Chisquare | df | *p* | Chisquare | df | *p* |
| Prokaryotic | Land-use Intensity (L) | 0.069 | 1 | 0.793 | 0.017 | 1 | 0.896 | 0.279 | 1 | 0.597 |
| | Grazing Intensity (G) | 0.066 | 1 | 0.797 | 0.029 | 1 | 0.865 | 0.036 | 1 | 0.850 |
| | Precipitation Treatment (P) | 1.773 | 4 | 0.777 | 1.860 | 4 | 0.762 | 1.215 | 4 | 0.876 |
| | L x G | 2.049 | 1 | 0.152 | 2.130 | 1 | 0.144 | 1.102 | 1 | 0.294 |
| | L x P | 2.996 | 4 | 0.559 | 2.401 | 4 | 0.662 | 3.553 | 4 | 0.470 |
| | G x P | 0.798 | 4 | 0.939 | 0.410 | 4 | 0.982 | 1.033 | 4 | 0.905 |
| | L x G x P | 3.351 | 4 | 0.501 | 3.287 | 4 | 0.511 | 3.441 | 4 | 0.487 |
| Fungi | Land-use Intensity (L) | 9.004 | 1 | **0.003** | 5.087 | 1 | **0.024** | 9.910 | 1 | **0.002** |
| | Grazing Intensity (G) | 0.948 | 1 | 0.330 | 2.075 | 1 | 0.150 | 0.884 | 1 | 0.347 |
| | Precipitation Treatment (P) | 4.002 | 4 | 0.406 | 4.994 | 4 | 0.288 | 5.223 | 4 | 0.265 |
| | L x G | 0.466 | 1 | 0.495 | 0.095 | 1 | 0.757 | 0.809 | 1 | 0.368 |
| | L x P | 7.196 | 4 | 0.126 | 3.275 | 4 | 0.513 | 8.457 | 4 | 0.076 |
| | G x P | 4.726 | 4 | 0.317 | 1.743 | 4 | 0.783 | 6.768 | 4 | 0.149 |
| | L x G x P | 4.130 | 4 | 0.389 | 1.852 | 4 | 0.763 | 4.758 | 4 | 0.313 |

**Table S2.** Results of PERMANOVA results showing how land-use intensity, grazing intensity, and altered precipitation treatments influence prokaryotic and fungal community composition. Bold values denote significant effects ($p < 0.05$).

| Community composition | Treatment | df | Sum of Sqs | Pseudo-*F* | *p* |
|---|---|---|---|---|---|
| Prokaryotic | Land-use Intensity (L) | 1 | 0.782 | 24.93 | **0.001** |
| | Grazing Intensity (G) | 1 | 0.034 | 1.07 | 0.320 |
| | Precipitation Treatment (P) | 4 | 0.104 | 0.83 | 0.635 |
| | L x G | 1 | 0.019 | 0.61 | 0.747 |
| | L x P | 4 | 0.104 | 0.83 | 0.634 |
| | G x P | 4 | 0.081 | 0.64 | 0.889 |
| | L x G x P | 4 | 0.139 | 1.11 | 0.229 |
| Fungi | Land-use Intensity (L) | 1 | 4.189 | 11.29 | **0.001** |
| | Grazing Intensity (G) | 1 | 0.378 | 1.02 | 0.274 |
| | Precipitation Treatment (P) | 4 | 1.252 | 0.84 | 0.768 |
| | L x G | 1 | 0.350 | 0.94 | 0.371 |
| | L x P | 4 | 1.342 | 0.9 | 0.561 |
| | G x P | 4 | 1.456 | 0.98 | 0.335 |
| | L x G x P | 4 | 1.300 | 0.88 | 0.643 |

**Table S3.** Prokaryotic and fungal taxa identified as important using random forest models with Boruta feature selection for distinguishing between land-use intensity.

| | Taxonomy (Phylum; Class; Order; Family) | Land-use intensity with higher abundance | Putative functional guild | Citation |
|---|---|---|---|---|
| Prokaryotic | Undefined | Intensively managed | — | — |
| | Euryarchaeota; Methanobacteria; Methanobacteriales; Methanobacteriaceae | Intensively managed | Hydrogenotrophic | Peng et al., 2024 |
| | Proteobacteria; Alphaproteobacteria; Rickettsiales; Mitochondria | Semi-natural | — | — |
| | Acidobacteriota; Acidobacteriae; Bryobacterales; Bryobacteraceae | Semi-natural | — | — |
| | Actinobacteriota; Acidimicrobiia; IMCC26256; IMCC26256 | Semi-natural | — | — |
| | Actinobacteriota; Actinobacteria; Frankiales; Acidothermaceae | Semi-natural | Saprotrophs, nutrient cyclers | Ma et al., 2025 |
| | Acidobacteriota; Acidobacteriae; Subgroup_2; Subgroup_2 | Semi-natural | — | — |
| | Proteobacteria; Alphaproteobacteria; Elsterales; uncultured | Semi-natural | — | — |
| Fungi | Ascomycota | Intensively managed | — | — |
| | Ascomycota; Archaeorhizomycetes; Archaeorhizomycetales; Archaeorhizomycetaceae | Intensively managed | Undefined Saprotroph | Rosling et al., (2011); Põlme et al., (2020) |
| | Ascomycota; Archaeorhizomycetes; Archaeorhizomycetales; Archaeorhizomycetaceae | Intensively managed | — | — |
| | Basidiomycota; Agaricomycetes; Agaricales; Mycenaceae | Intensively managed | Plant Saprotroph | Tedersoo et al., (2014); Põlme et al., (2020) |

| | Taxonomy (Phylum; Class; Order; Family) | Land-use intensity with higher abundance | Putative functional guild | Citation |
|---|---|---|---|---|
| Fungi | Basidiomycota; Agaricomycetes; Agaricales | Intensively managed | — | — |
| | Basidiomycota; Agaricomycetes | Intensively managed | — | — |
| | Rozellomycota; Rozellomycotina_cls_undefined; GS11; GS11_fam_undefined | Intensively managed | — | — |
| | Ascomycota; Leotiomycetes; Helotiales | Semi-natural | — | — |
| | Ascomycota; Sordariomycetes; Chaetosphaeriales; Chaetosphaeriaceae | Semi-natural | Plant Saprotroph-Wood Saprotroph | Cannon and Kirk, (2007) |
| | Undefined | Semi-natural | — | — |
| | Ascomycota; Sordariomycetes | Semi-natural | — | — |
| | Ascomycota; Sordariomycetes; Hypocreales; Ophiocordycipitaceae | Semi-natural | — | — |
| | Ascomycota; Sordariomycetes; Sordariales; Lasiosphaeriaceae | Semi-natural | Dung Saprotroph | Bills et al., (2013); Tedersoo et al., (2014;) Põlme et al., (2020) |
| | Ascomycota; Sordariomycetes; Amphisphaeriales; Sporocadaceae | Semi-natural | Plant Pathogen | Põlme et al., (2020) |
| | Ascomycota; Dothideomycetes; Venturiales; Venturiales_fam_undefined | Semi-natural | — | — |
| | Chytridiomycota; Chytridiomycota_cls_undefined; Chytridiomycota_ord_undefined; Chytridiomycota_fam_undefined | Semi-natural | — | — |
| | Ascomycota; Pezizomycetes; Pezizales; Pyronemataceae | Semi-natural | Undefined Saprotroph | Tedersoo et al., (2014); Põlme et al., (2020) |

| | Taxonomy (Phylum; Class; Order; Family) | Land-use intensity with higher abundance | Putative functional guild | Citation |
|---|---|---|---|---|
| | Ascomycota; Eurotiomycetes; Chaetothyriales; Herpotrichiellaceae | Semi-natural | Undefined Saprotroph | James et al., (2006); Põlme, et al., (2020) |
| | Entorrhizomycota; Entorrhizomycetes; Entorrhizales; Entorrhizaceae | Semi-natural | Plant Pathogen | Tedersoo et al., (2014); Põlme et al., (2020) |
| | Ascomycota; Dothideomycetes; Jahnulales; Aliquandostipitaceae | Semi-natural | Wood Saprotroph | Tedersoo et al., (2014); Põlme et al., (2020) |
| | Ascomycota; Ascomycota_cls_undefined; Ascomycota_ord_undefined; Ascomycota_fam_undefined | Semi-natural | — | — |
| | Ascomycota; Dothideomycetes; Pleosporales; Didymosphaeriaceae | Semi-natural | — | — |
| Fungi | Ascomycota; Eurotiomycetes; Eurotiales; Aspergillaceae | Semi-natural | — | — |
| | Ascomycota; Eurotiomycetes; Chaetothyriales; Herpotrichiellaceae | Semi-natural | — | — |
| | Ascomycota; Saccharomycetes; Saccharomycetales; Lipomycetaceae | Semi-natural | — | — |
| | Ascomycota; Eurotiomycetes; Chaetothyriales; Herpotrichiellaceae | Semi-natural | Animal Pathogen-Fungal Parasite-Undefined Saprotroph | Cannon and Kirk, (2007) |
| | Ascomycota; Dothideomycetes; Dothideomycetes_ord_undefined; Dothideomycetes_fam_undefined | Semi-natural | — | — |
| | Ascomycota; Dothideomycetes; Pleosporales; Sporormiaceae | Semi-natural | Dung Saprotroph | Tedersoo et al., (2014); Põlme et al., (2020) |

| Taxonomy (Phylum; Class; Order; Family) | Land-use intensity with higher abundance | Putative functional guild | Citation |
|---|---|---|---|
| Mortierellomycota; Mortierellomycetes; Mortierellales; Mortierellaceae | Semi-natural | — | — |
| Basidiomycota | Semi-natural | — | — |

Note: Columns with '—' indicate unidentified. Putative functional guilds were determined using literature searches and FUNGuild with corresponding citations provided.

**Table S4.** Fungal taxa identified as important using random forest models with Boruta feature selection for distinguishing between grazing intensity.

| Taxonomy (Phylum; Class; Order; Family) | Grazing intensity with higher abundance | Putative functional guild | Citation |
|---|---|---|---|
| Ascomycota; Dothideomycetes; Pleosporales; Didymosphaeriaceae | Low grazing | Wood Saprotroph | Tedersoo et al., (2014) ; Põlme et al., (2020) |
| Basidiomycota; Agaricomycetes; —; — | Low grazing | — | — |

Note: Columns with '—' indicate unidentified. Putative functional guilds and citations were determined using FUNGuild.

**Table S5.** Prokaryotic and fungal taxa identified as important using random forest models with Boruta feature selection for distinguishing between precipitation treatments.

| | Taxonomy (Phylum; Class; Order; Family) | Precipitation treatment with higher abundance | Putative functional guild | Citation |
|---|---|---|---|---|
| Prokaryotic | Firmicutes; Bacilli; Bacillales; Bacillaceae | Diminished seasonality | Decomposers of organic matter content, plant health and growth | Mandic-Mulec et al., 2015 |
| Fungi | Ascomycota; Eurotiomycetes; Chaetothyriales; — | Control | — | — |

Note: Columns with '—' indicate unidentified. Putative functional guilds were determined using literature searches and FUNGuild with corresponding citations provided.

**Table S6.** PERMANOVA results showing how land-use intensity, grazing intensity, and precipitation treatments influence prokaryotic functional KEGG pathway composition. Bold values denote significant effects ($p < 0.05$).

| KEGG Pathways | Treatment | df | Sum of Sqs | Pseudo-$F$ | $p$ |
|---|---|---|---|---|---|
| Prokaryotic | Land-use Intensity (L) | 1 | 0.016 | 23.35 | **0.001** |
| | Grazing Intensity (G) | 1 | 0.001 | 1.01 | 0.353 |
| | Precipitation Treatment (P) | 4 | 0.003 | 0.93 | 0.472 |
| | L x G | 1 | 0.000 | 0.65 | 0.649 |
| | L x P | 4 | 0.002 | 0.78 | 0.642 |
| | G x P | 4 | 0.003 | 1.03 | 0.323 |
| | L x G x P | 4 | 0.003 | 0.95 | 0.430 |

**Table S7.** Results of generalized linear mixed models of land-use intensity, grazing intensity, and precipitation treatment effects on the three most common fungal functional guilds (undefined saprotroph, wood saprotroph, and plant saprotroph). Bold values denote significant effects ($p$ < 0.05).

| | Treatment | Chisq | df | *p* |
|---|---|---|---|---|
| Guild Richness | Land-use Intensity (L) | 4.226 | 1 | **0.040** |
| | Grazing Intensity (G) | 1.056 | 1 | 0.304 |
| | Precipitation Treatment (P) | 3.357 | 4 | 0.500 |
| | L x G | 0.015 | 1 | 0.904 |
| | L x P | 0.998 | 4 | 0.910 |
| | G x P | 2.899 | 4 | 0.575 |
| | L x G x P | 1.379 | 4 | 0.848 |
| Undefined Saprotroph | Land-use Intensity (L) | 3.935 | 1 | **0.047** |
| | Grazing Intensity (G) | 0.653 | 1 | 0.419 |
| | Precipitation Treatment (P) | 6.443 | 4 | 0.168 |
| | L x G | 0.151 | 1 | 0.697 |
| | L x P | 8.326 | 4 | 0.080 |
| | G x P | 1.387 | 4 | 0.847 |
| | L x G x P | 2.051 | 4 | 0.726 |
| Wood Saprotroph | Land-use Intensity (L) | 1.178 | 1 | 0.278 |
| | Grazing Intensity (G) | 0.330 | 1 | 0.566 |
| | Precipitation Treatment (P) | 1.613 | 4 | 0.806 |
| | L x G | 0.264 | 1 | 0.607 |
| | L x P | 12.754 | 4 | **0.013** |
| | G x P | 3.221 | 4 | 0.522 |
| | L x G x P | 5.440 | 4 | 0.245 |
| Plant Saprotroph | Land-use Intensity (L) | 6.660 | 1 | **0.010** |
| | Grazing Intensity (G) | 0.069 | 1 | 0.793 |
| | Precipitation Treatment (P) | 3.117 | 4 | 0.538 |
| | L x G | 0.809 | 1 | 0.368 |
| | L x P | 9.959 | 4 | **0.041** |
| | G x P | 3.102 | 4 | 0.541 |
| | L x G x P | 2.290 | 4 | 0.683 |

**Table S8.** Network metrics modularity, negative:positive cohesion ratio, and richness for prokaryotes and fungi. Samples from the same treatment combination of land-use intensity and precipitation were combined to construct one microbial network for each microbial group (prokaryotes and fungi).

| | Land-use Intensity | Precipitation Treatment | Modularity | Negative:positive cohesion | Richness |
|---|---|---|---|---|---|
| Prokaryotes | Semi-natural | Increased quantity | 0.629 | 0.428 | 103 |
| | | Reduced quantity | 0.600 | 0.702 | 90 |
| | | Enhanced seasonality | 0.512 | 0.800 | 122 |
| | | Diminished seasonality | 0.305 | 0.574 | 87 |
| | | Control | 0.518 | 0.768 | 120 |
| | Intensively managed | Increased quantity | 0.644 | 0.564 | 48 |
| | | Reduced quantity | 0.675 | 1.003 | 38 |
| | | Enhanced seasonality | 0.692 | 0.389 | 45 |
| | | Diminished seasonality | 0.752 | 0.503 | 59 |
| | | Control | 0.659 | 0.395 | 38 |
| Fungi | Semi-natural | Increased quantity | 0.475 | 0.961 | 143 |
| | | Reduced quantity | 0.429 | 1.112 | 134 |
| | | Enhanced seasonality | 0.476 | 0.920 | 146 |
| | | Diminished seasonality | 0.509 | 0.977 | 149 |
| | | Control | 0.454 | 0.967 | 153 |
| | Intensively managed | Increased quantity | 0.506 | 0.833 | 106 |
| | | Reduced quantity | 0.517 | 0.945 | 107 |
| | | Enhanced seasonality | 0.488 | 0.701 | 131 |
| | | Diminished seasonality | 0.487 | 0.930 | 118 |
| | | Control | 0.530 | 0.917 | 85 |

**Figure S1.**

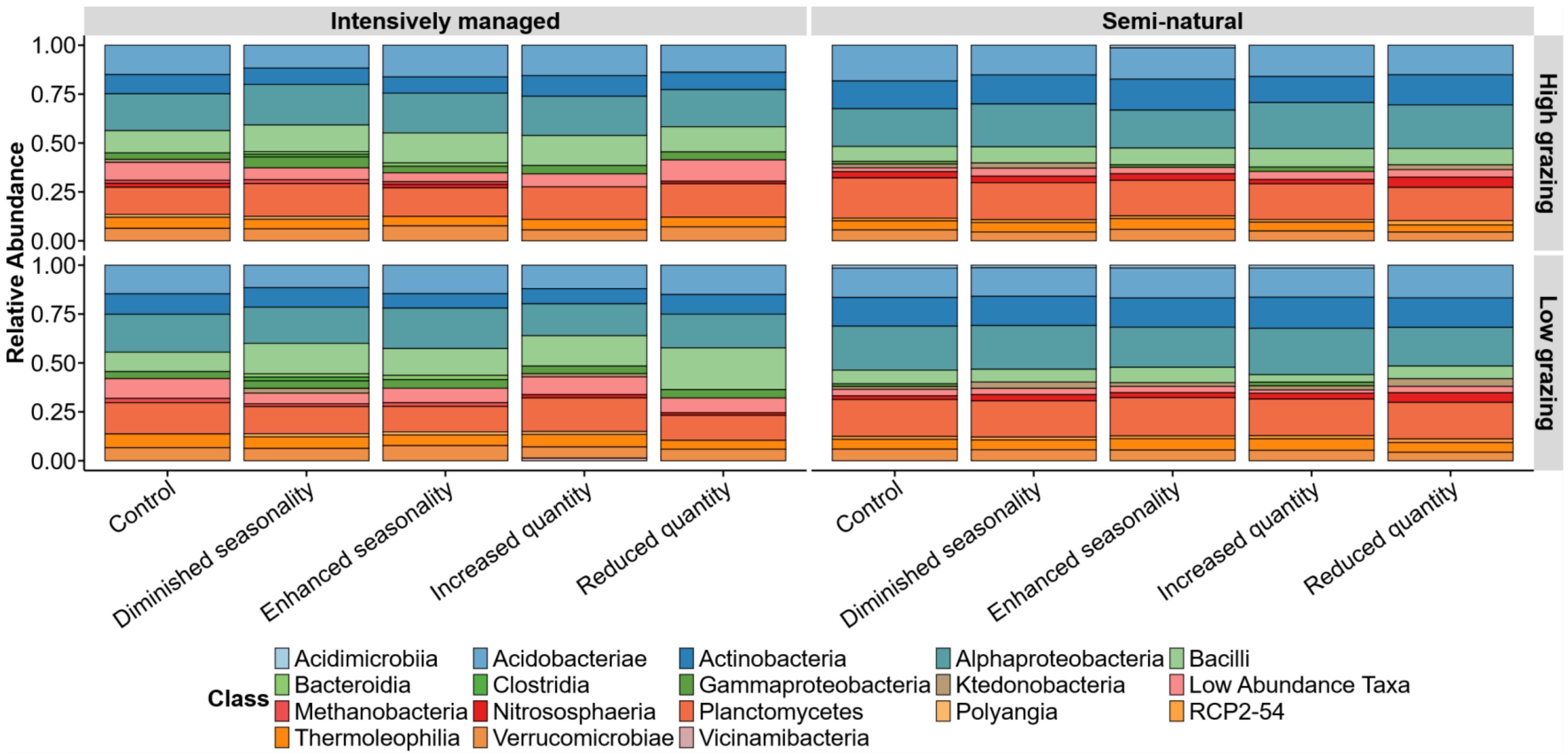


**Figure S1. Prokaryotic relative abundance at the class level across experimental treatments.** Each stacked bar represents a unique combination of land-use intensity, grazing intensity, and precipitation treatments. Microbial classes with a relative abundance of less than 0.05% across all samples were aggregated into a “low abundance” category.

**Figure S2.**

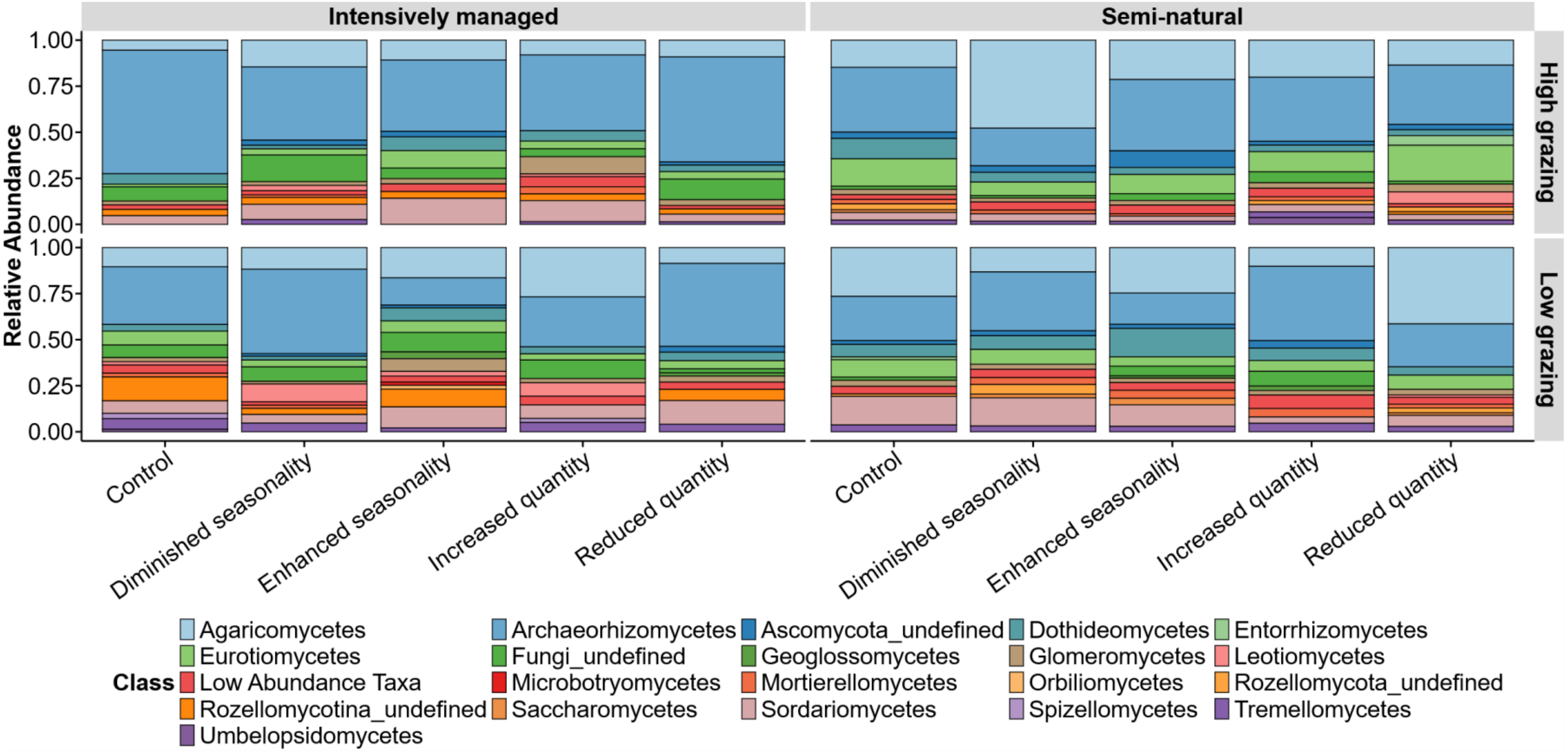


**Figure S2. Fungal relative abundance at the class level across experimental treatments.** Each stacked bar represents a unique combination of land-use intensity, grazing intensity, and precipitation treatments. Microbial classes with a relative abundance of less than 0.05% across all samples were aggregated into a “low abundance” category.

**Figure S3.**

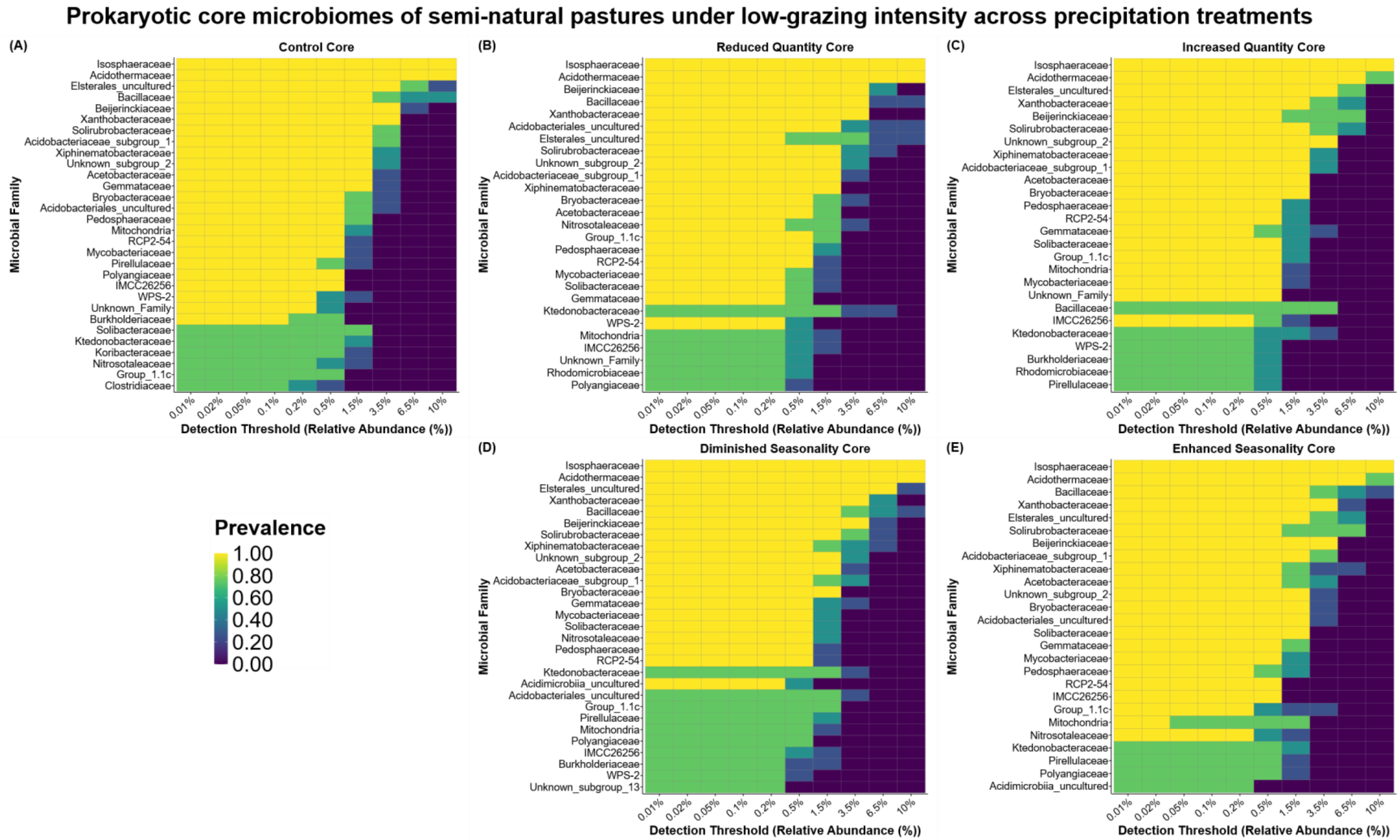


**Figure S3. Core prokaryotic microbial families in semi-natural pastures under low-intensity grazing across precipitation treatments.** Microbial families are organized by relative abundance detection thresholds and shaded by prevalence.

**Figure S4.**

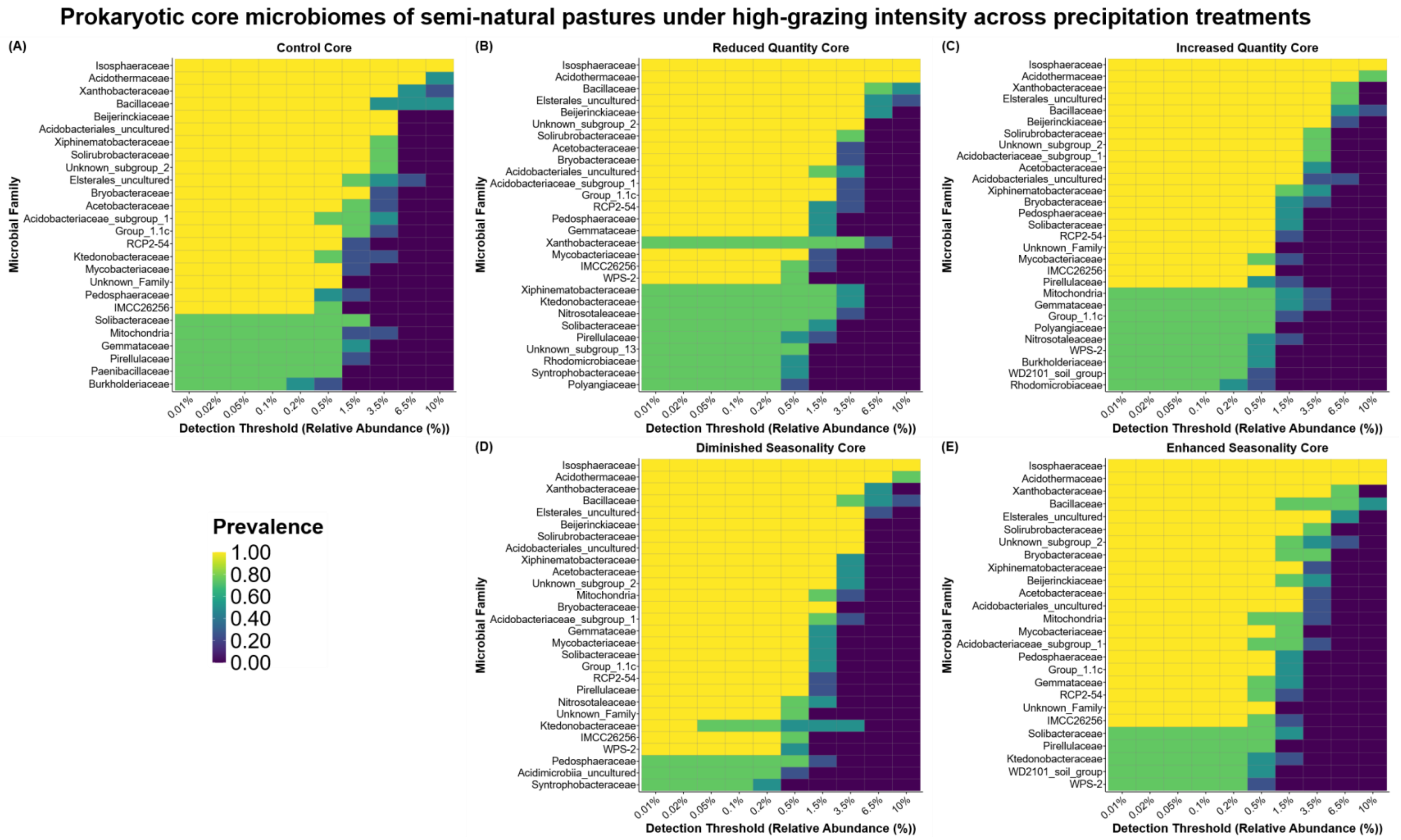


**Figure S4. Core prokaryotic microbial families in semi-natural pastures under high-intensity grazing across precipitation treatments.** Microbial families are organized by relative abundance detection thresholds and shaded by prevalence.

**Figure S5.**

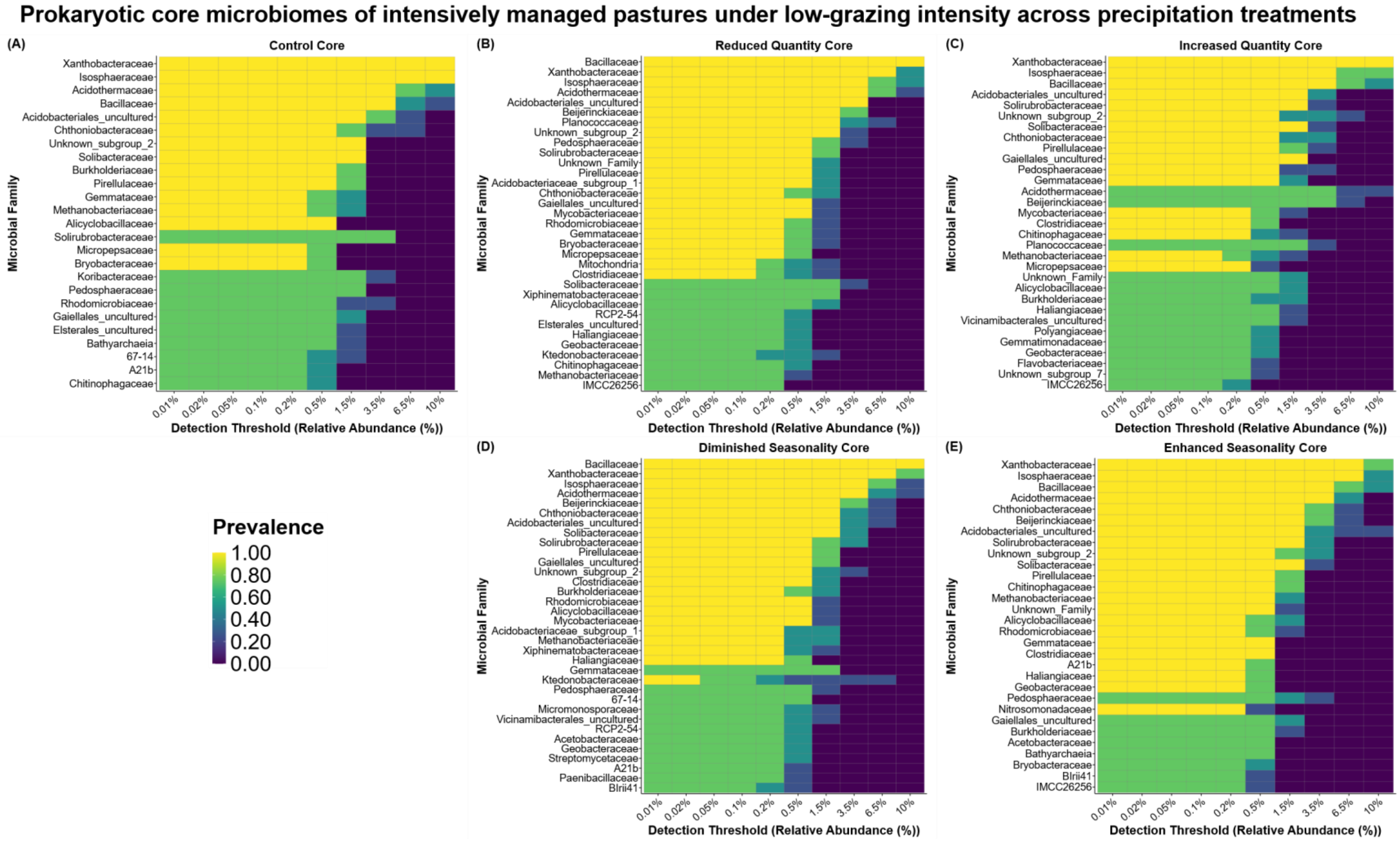


**Figure S5. Core prokaryotic microbial families in intensively managed pastures under low-intensity grazing across precipitation treatments.** Microbial families are organized by relative abundance detection thresholds and shaded by prevalence.

**Figure S6.**

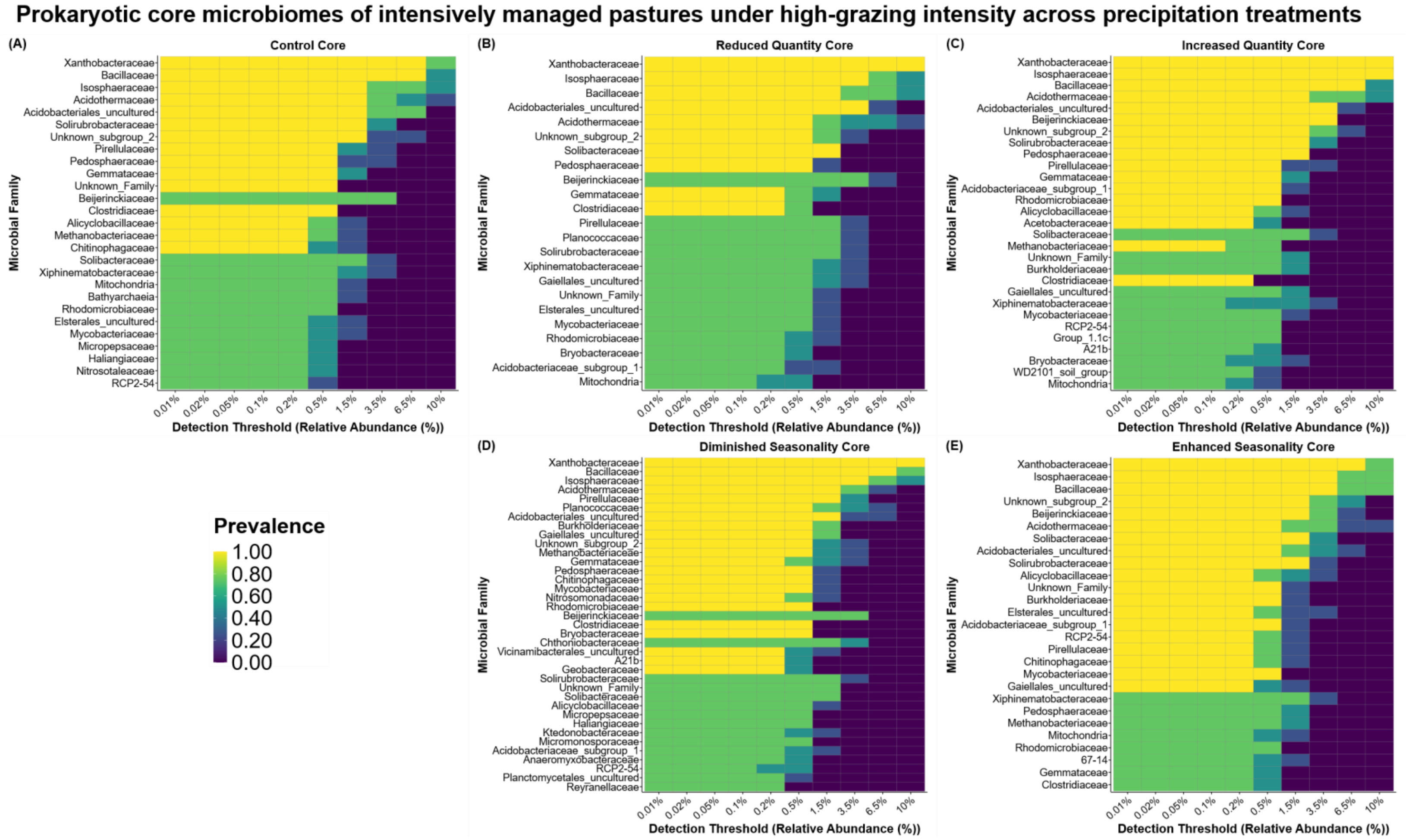


**Figure S6. Core prokaryotic microbial families in intensively managed pastures under high-intensity grazing across precipitation treatments.** Microbial families are organized by relative abundance detection thresholds and shaded by prevalence.

**Figure S7.**

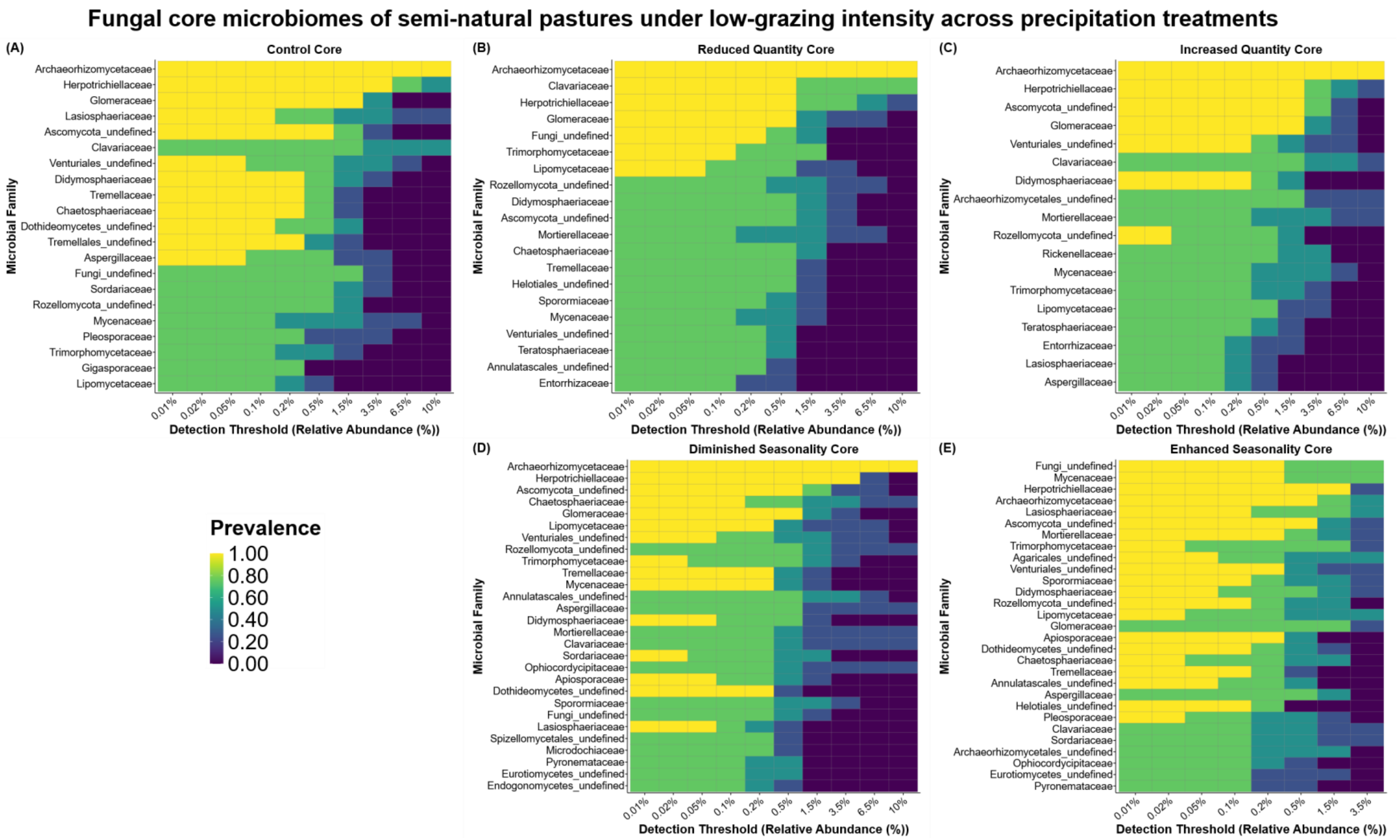


**Figure S7. Core fungal microbial families in semi-natural pastures under low-intensity grazing across precipitation treatments.** Microbial families are organized by relative abundance detection thresholds and shaded by prevalence.

**Figure S8.**

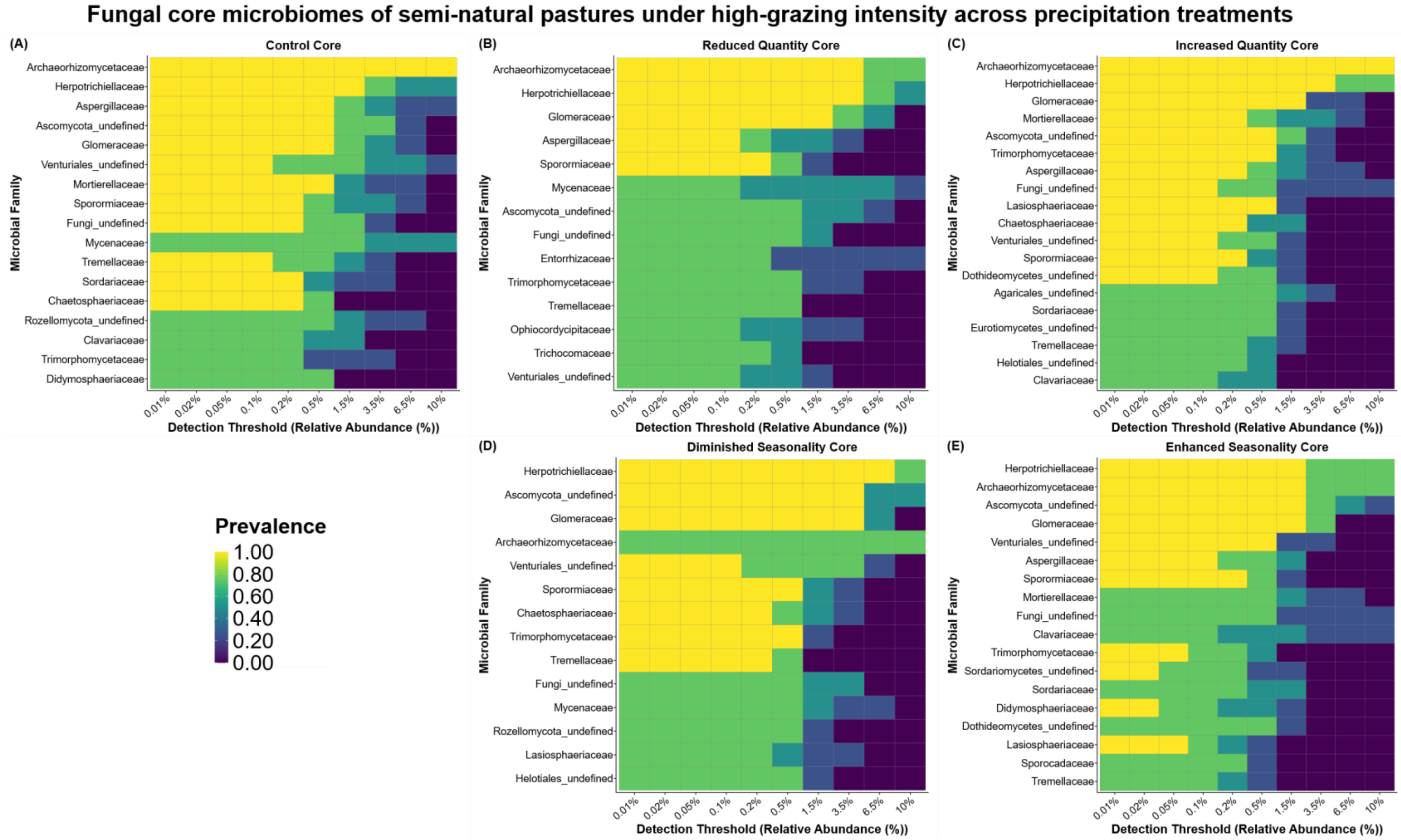


**Figure S8. Core fungal microbial families in semi-natural pastures under high-intensity grazing across precipitation treatments.** Microbial families are organized by relative abundance detection thresholds and shaded by prevalence.

75

**Figure S9.**

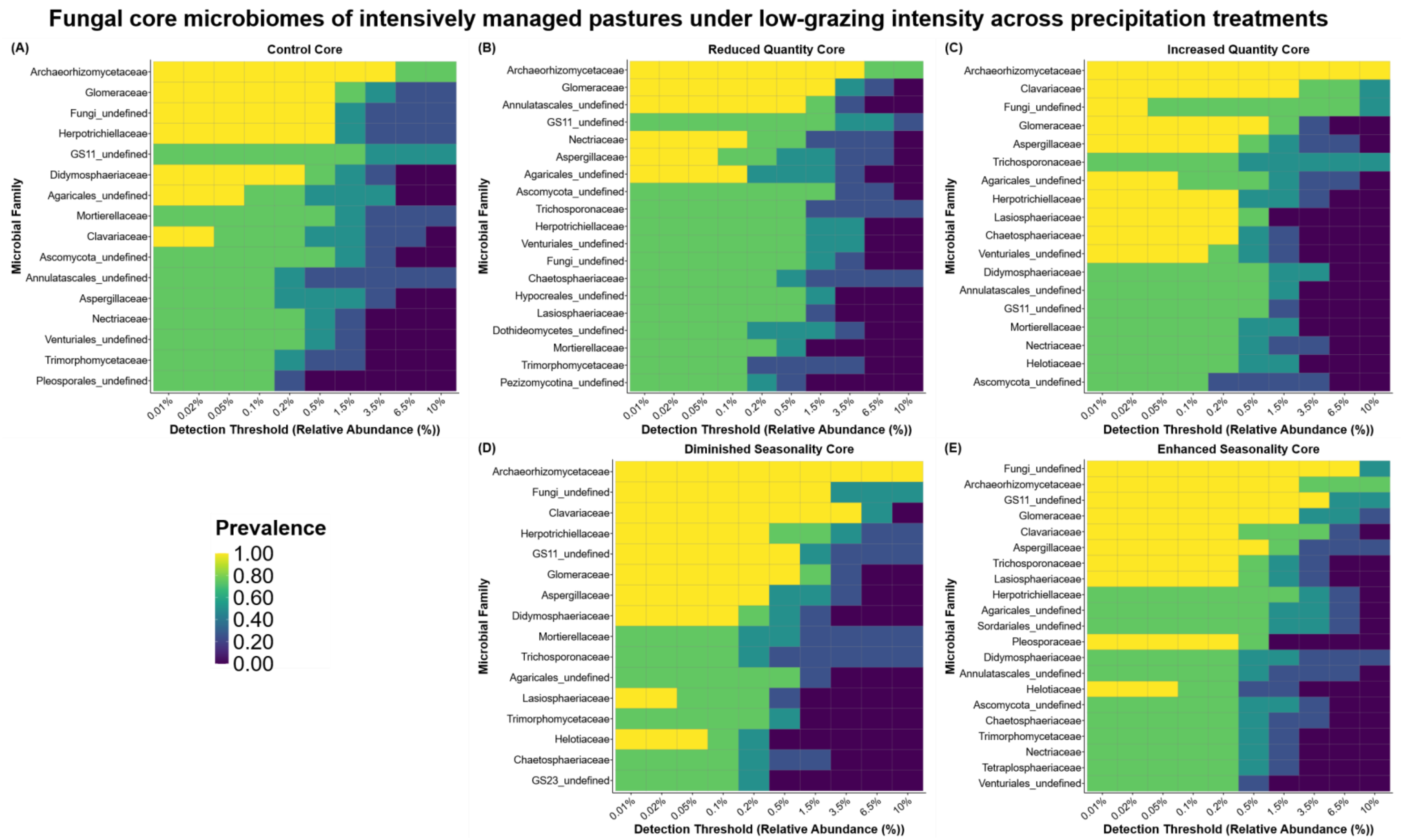
Fungal core microbiomes of intensively managed pastures under low-grazing intensity across precipitation treatments
(A)
Control Core
Archaeorhizomycetaceae
Glomeraceae
Fungi_undefined
Herpotrichiellaceae
GS11_undefined
Didymosphaeriaceae
Agaricales_undefined
Mortierellaceae
Clavariaceae
Ascomycota_undefined
Annulatascales_undefined
Aspergillaceae
Nectriaceae
Venturiales_undefined
Trimorphomycetaceae
Pleosporales_undefined
Microbial Family
0.01% 0.02% 0.05% 0.1% 0.2% 0.5% 1.5% 3.5% 6.5% 10%
Detection Threshold (Relative Abundance (%))
(B)
Reduced Quantity Core
Archaeorhizomycetaceae
Glomeraceae
Annulatascales_undefined
GS11_undefined
Nectriaceae
Aspergillaceae
Agaricales_undefined
Ascomycota_undefined
Trichosporonaceae
Herpotrichiellaceae
Venturiales_undefined
Fungi_undefined
Chaetosphaeriaceae
Hypocreales_undefined
Lasiosphaeriaceae
Dothideomycetes_undefined
Mortierellaceae
Trimorphomycetaceae
Pezizomycotina_undefined
Microbial Family
0.01% 0.02% 0.05% 0.1% 0.2% 0.5% 1.5% 3.5% 6.5% 10%
Detection Threshold (Relative Abundance (%))
(C)
Increased Quantity Core
Archaeorhizomycetaceae
Clavariaceae
Fungi_undefined
Glomeraceae
Aspergillaceae
Trichosporonaceae
Agaricales_undefined
Herpotrichiellaceae
Lasiosphaeriaceae
Chaetosphaeriaceae
Venturiales_undefined
Didymosphaeriaceae
Annulatascales_undefined
GS11_undefined
Mortierellaceae
Nectriaceae
Helotiaceae
Ascomycota_undefined
Microbial Family
0.01% 0.02% 0.05% 0.1% 0.2% 0.5% 1.5% 3.5% 6.5% 10%
Detection Threshold (Relative Abundance (%))
Prevalence
1.00
0.80
0.60
0.40
0.20
0.00
(D)
Diminished Seasonality Core
Archaeorhizomycetaceae
Fungi_undefined
Clavariaceae
Herpotrichiellaceae
GS11_undefined
Glomeraceae
Aspergillaceae
Didymosphaeriaceae
Mortierellaceae
Trichosporonaceae
Agaricales_undefined
Lasiosphaeriaceae
Trimorphomycetaceae
Helotiaceae
Chaetosphaeriaceae
GS23_undefined
Microbial Family
0.01% 0.02% 0.05% 0.1% 0.2% 0.5% 1.5% 3.5% 6.5% 10%
Detection Threshold (Relative Abundance (%))
(E)
Enhanced Seasonality Core
Fungi_undefined
Archaeorhizomycetaceae
GS11_undefined
Glomeraceae
Clavariaceae
Aspergillaceae
Trichosporonaceae
Lasiosphaeriaceae
Herpotrichiellaceae
Agaricales_undefined
Sordariales_undefined
Pleosporaceae
Didymosphaeriaceae
Annulatascales_undefined
Helotiaceae
Ascomycota_undefined
Chaetosphaeriaceae
Trimorphomycetaceae
Nectriaceae
Tetraplosphaeriaceae
Venturiales_undefined
Microbial Family
0.01% 0.02% 0.05% 0.1% 0.2% 0.5% 1.5% 3.5% 6.5% 10%
Detection Threshold (Relative Abundance (%))

**Figure S9. Core fungal microbial families in intensively managed pastures under low-intensity grazing across precipitation treatments.** Microbial families are organized by relative abundance detection thresholds and shaded by prevalence.

**Figure S10.**

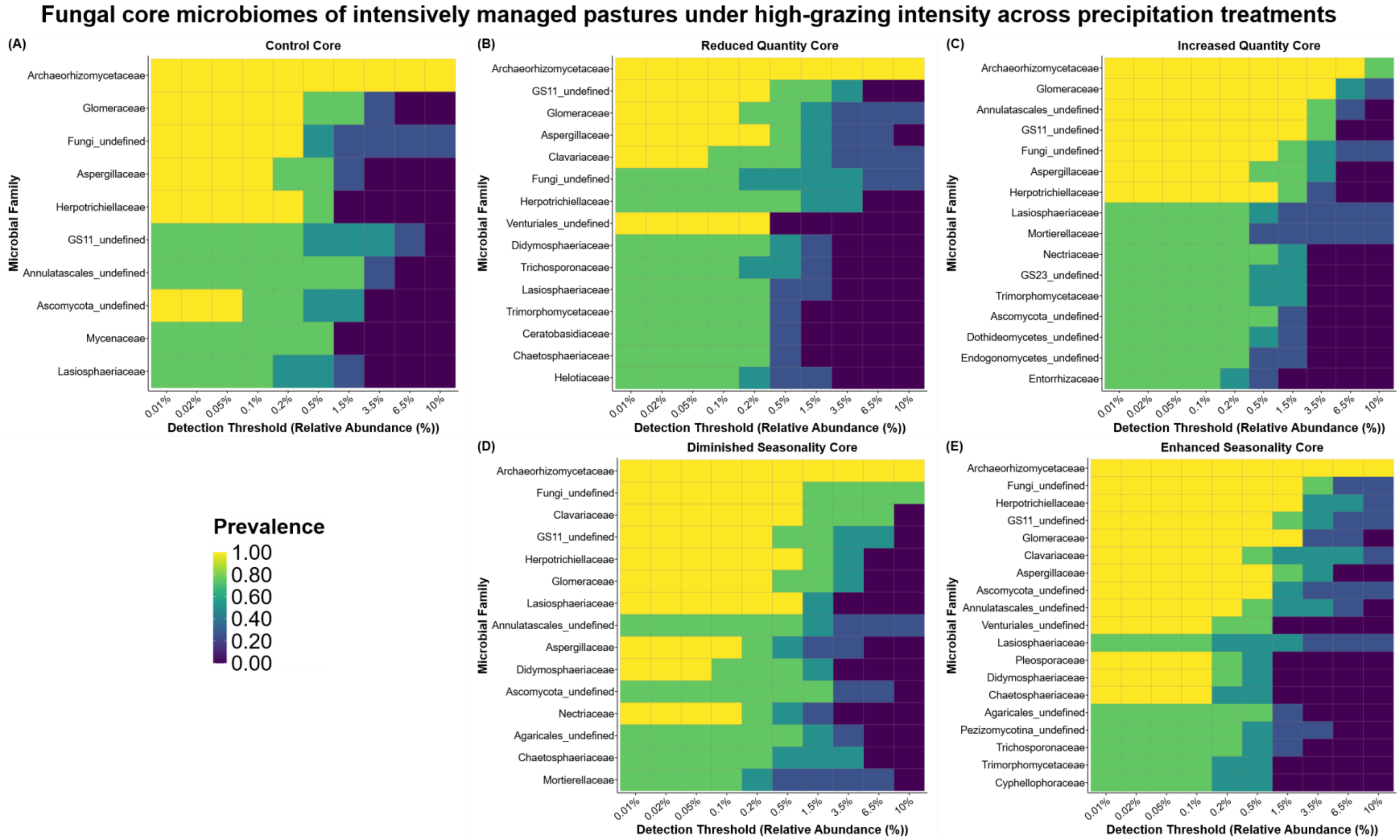


**Figure S10. Core fungal microbial families in intensively managed pastures under high-intensity grazing across precipitation treatments.** Microbial families are organized by relative abundance detection thresholds and shaded by prevalence.

**Figure S11.**

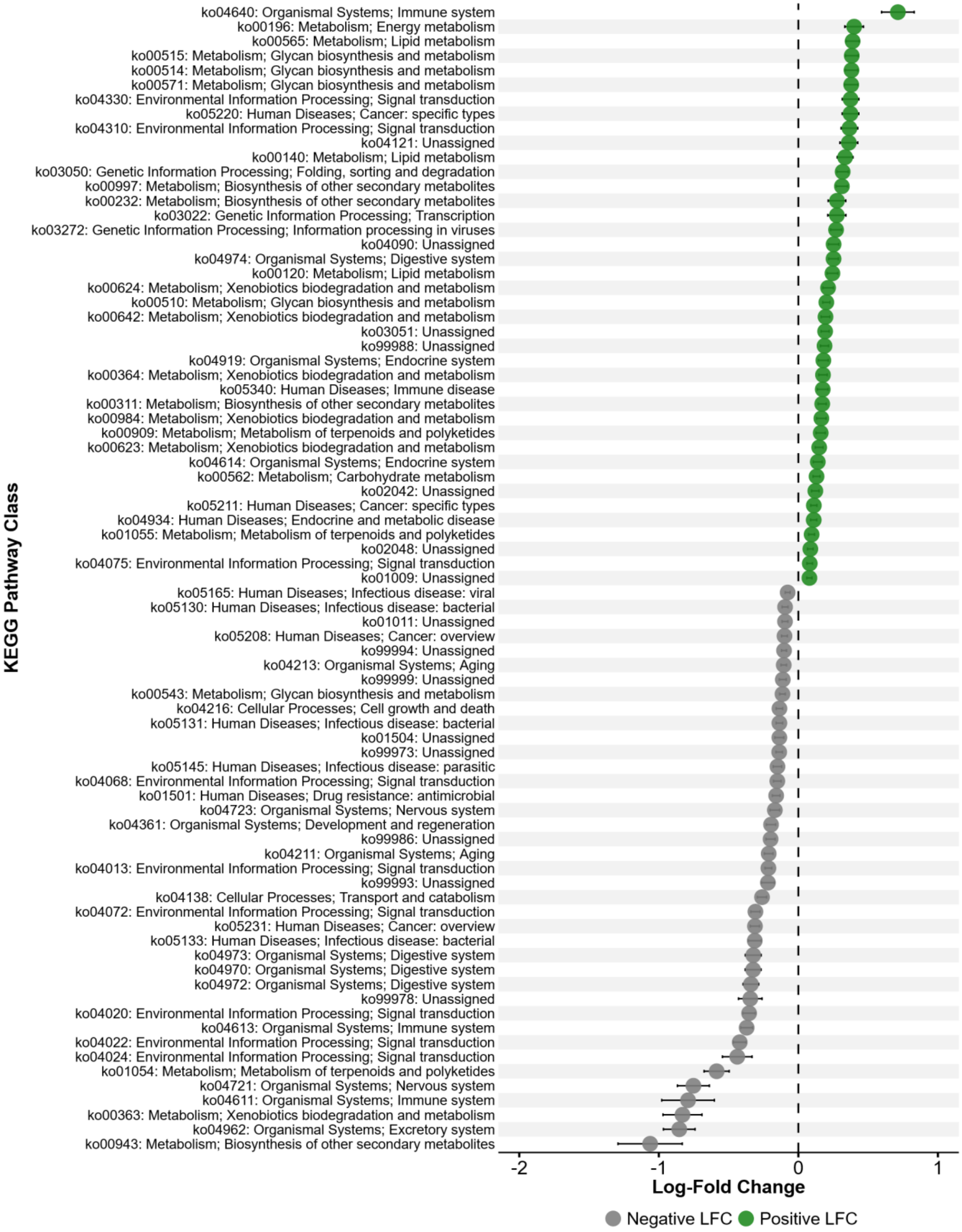
ko04640: Organismal Systems; Immune system
ko00196: Metabolism; Energy metabolism
ko00565: Metabolism; Lipid metabolism
ko00515: Metabolism; Glycan biosynthesis and metabolism
ko00514: Metabolism; Glycan biosynthesis and metabolism
ko00571: Metabolism; Glycan biosynthesis and metabolism
ko04330: Environmental Information Processing; Signal transduction
ko05220: Human Diseases; Cancer: specific types
ko04310: Environmental Information Processing; Signal transduction
ko04121: Unassigned
ko00140: Metabolism; Lipid metabolism
ko03050: Genetic Information Processing; Folding, sorting and degradation
ko00997: Metabolism; Biosynthesis of other secondary metabolites
ko00232: Metabolism; Biosynthesis of other secondary metabolites
ko03022: Genetic Information Processing; Transcription
ko03272: Genetic Information Processing; Information processing in viruses
ko04090: Unassigned
ko04974: Organismal Systems; Digestive system
ko00120: Metabolism; Lipid metabolism
ko00624: Metabolism; Xenobiotics biodegradation and metabolism
ko00510: Metabolism; Glycan biosynthesis and metabolism
ko00642: Metabolism; Xenobiotics biodegradation and metabolism
ko03051: Unassigned
ko99988: Unassigned
ko04919: Organismal Systems; Endocrine system
ko00364: Metabolism; Xenobiotics biodegradation and metabolism
ko05340: Human Diseases; Immune disease
ko00311: Metabolism; Biosynthesis of other secondary metabolites
ko00984: Metabolism; Xenobiotics biodegradation and metabolism
ko00909: Metabolism; Metabolism of terpenoids and polyketides
ko00623: Metabolism; Xenobiotics biodegradation and metabolism
ko04614: Organismal Systems; Endocrine system
ko00562: Metabolism; Carbohydrate metabolism
ko02042: Unassigned
ko05211: Human Diseases; Cancer: specific types
ko04934: Human Diseases; Endocrine and metabolic disease
ko01055: Metabolism; Metabolism of terpenoids and polyketides
ko02048: Unassigned
ko04075: Environmental Information Processing; Signal transduction
ko01009: Unassigned
ko05165: Human Diseases; Infectious disease: viral
ko05130: Human Diseases; Infectious disease: bacterial
ko01011: Unassigned
ko05208: Human Diseases; Cancer: overview
ko99994: Unassigned
ko04213: Organismal Systems; Aging
ko99999: Unassigned
ko00543: Metabolism; Glycan biosynthesis and metabolism
ko04216: Cellular Processes; Cell growth and death
ko05131: Human Diseases; Infectious disease: bacterial
ko01504: Unassigned
ko99973: Unassigned
ko05145: Human Diseases; Infectious disease: parasitic
ko04068: Environmental Information Processing; Signal transduction
ko01501: Human Diseases; Drug resistance: antimicrobial
ko04723: Organismal Systems; Nervous system
ko04361: Organismal Systems; Development and regeneration
ko99986: Unassigned
ko04211: Organismal Systems; Aging
ko04013: Environmental Information Processing; Signal transduction
ko99993: Unassigned
ko04138: Cellular Processes; Transport and catabolism
ko04072: Environmental Information Processing; Signal transduction
ko05231: Human Diseases; Cancer: overview
ko05133: Human Diseases; Infectious disease: bacterial
ko04973: Organismal Systems; Digestive system
ko04970: Organismal Systems; Digestive system
ko04972: Organismal Systems; Digestive system
ko99978: Unassigned
ko04020: Environmental Information Processing; Signal transduction
ko04613: Organismal Systems; Immune system
ko04022: Environmental Information Processing; Signal transduction
ko04024: Environmental Information Processing; Signal transduction
ko01054: Metabolism; Metabolism of terpenoids and polyketides
ko04721: Organismal Systems; Nervous system
ko04611: Organismal Systems; Immune system
ko00363: Metabolism; Xenobiotics biodegradation and metabolism
ko04962: Organismal Systems; Excretory system
ko00943: Metabolism; Biosynthesis of other secondary metabolites
KEGG Pathway Class
-2
-1
0
1
Log-Fold Change
Negative LFC
Positive LFC

**Figure S11. Land-use intensity drives differences in prokaryotic functional KEGG pathways between intensively managed and semi-natural pastures.** Differentially abundant KEGG pathway classes identified by ANCOM-BC2. All significant pathways in the land-use intensity comparison are shown. Intensively managed pasture was used as the reference level. Positive values indicate higher abundance in semi-natural pastures. Colors indicate positive or negative log-fold change (LFC) of KEGG pathway classes relative to the reference. Points represent mean LFC and error bars represent SE from ANCOM-BC2.

**Figure S12.**

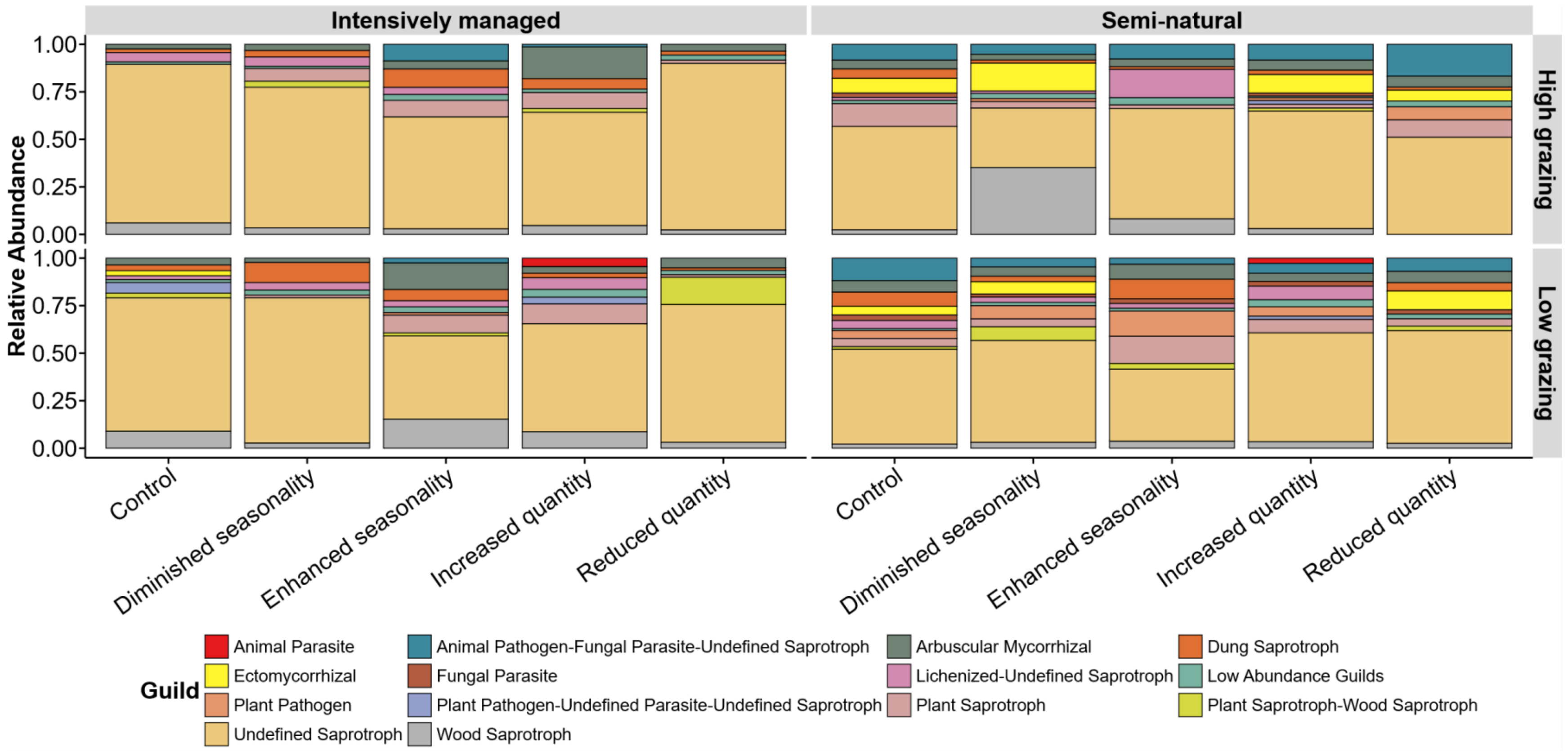


**Figure S12. Fungal functional guild relative abundance across experimental treatments.** Each stacked bar represents a unique combination of land-use intensity, grazing intensity, and precipitation treatments. Fungal functional guilds with a relative abundance of less than 0.05% across all samples were aggregated into a "low abundance" category.

**Supplementary References – Reyes et al., 2026 – Land-use intensity overrides grazing and precipitation effects on soil microbial communities in a subtropical agroecosystem**


Bills, G.F., Gloer, J.B., An, Z., 2013. Coprophilous fungi: antibiotic discovery and functions in an underexplored arena of microbial defensive mutualism. Current Opinion in Microbiology 16, 549–565. https://doi.org/10.1016/j.mib.2013.08.001

Cannon, P.F., Kirk, P.M., 2007. Fungal Families of the World. CABI.

James, T.Y., Kauff, F., Schoch, C.L., Matheny, P.B., Hofstetter, V., Cox, C.J., Celio, G., Gueidan, C., Fraker, E., Miadlikowska, J., Lumbsch, H.T., Rauhut, A., Reeb, V., Arnold, A.E., Amtoft, A., Stajich, J.E., Hosaka, K., Sung, G.-H., Johnson, D., O'Rourke, B., Crockett, M., Binder, M., Curtis, J.M., Slot, J.C., Wang, Z., Wilson, A.W., Schüßler, A., Longcore, J.E., O'Donnell, K., Mozley-Standridge, S., Porter, D., Letcher, P.M., Powell, M.J., Taylor, J.W., White, M.M., Griffith, G.W., Davies, D.R., Humber, R.A., Morton, J.B., Sugiyama, J., Rossman, A.Y., Rogers, J.D., Pfister, D.H., Hewitt, D., Hansen, K., Hambleton, S., Shoemaker, R.A., Kohlmeyer, J., Volkmann-Kohlmeyer, B., Spotts, R.A., Serdani, M., Crous, P.W., Hughes, K.W., Matsuura, K., Langer, E., Langer, G., Untereiner, W.A., Lücking, R., Büdel, B., Geiser, D.M., Aptroot, A., Diederich, P., Schmitt, I., Schultz, M., Yahr, R., Hibbett, D.S., Lutzoni, F., McLaughlin, D.J., Spatafora, J.W., Vilgalys, R., 2006. Reconstructing the early evolution of Fungi using a six-gene phylogeny. Nature 443, 818–822. https://doi.org/10.1038/nature05110

Ma, J., Liu, W., Wang, M., Ye, Z., Liu, D., 2025. Lei bamboo (Phyllostachys praecox) shows greater sensitivity to salt stress than to hypoxia stress: insights from plant physiology, metabolome and soil microbiome. Plant Soil 513, 2395–2415. https://doi.org/10.1007/s11104-025-07322-9

Mandic-Mulec, I., Stefanic, P., Van Elsas, J.D., 2015. Ecology of Bacillaceae. Microbiol Spectr 3, 3.2.16. https://doi.org/10.1128/microbiolspec.TBS-0017-2013

Peng, W., Lu, J., Kuang, J., Tang, R., Guan, F., Xie, K., Zhou, L., Yuan, Y., 2024. Enhancement of hydrogenotrophic methanogenesis for methane production by nano zero-valent iron in soils. Environmental Research 247, 118232. https://doi.org/10.1016/j.envres.2024.118232

Põlme, S., Abarenkov, K., Henrik Nilsson, R., Lindahl, B.D., Clemmensen, K.E., Kauserud, H., Nguyen, N., Kjøller, R., Bates, S.T., Baldrian, P., Frøslev, T.G., Adojaan, K., Vizzini, A., Suija, A., Pfister, D., Baral, H.-O., Järv, H., Madrid, H., Nordén, J., Liu, J.-K., Pawlowska, J., Põldmaa, K., Pärtel, K., Runnel, K., Hansen, K., Larsson, K.-H., Hyde, K.D., Sandoval-Denis, M., Smith, M.E., Toome-Heller, M., Wijayawardene, N.N., Menolli, N., Reynolds, N.K., Drenkhan, R., Maharachchikumbura, S.S.N., Gibertoni, T.B., Læssøe, T., Davis, W., Tokarev, Y., Corrales, A., Soares, A.M., Agan, A., Machado, A.R., Argüelles-Moyao, A., Detheridge, A., de Meiras-Ottoni, A., Verbeken, A., Dutta, A.K., Cui, B.-K., Pradeep, C.K., Marín, C., Stanton, D., Gohar, D., Wanasinghe, D.N., Otsing, E., Aslani, F., Griffith, G.W., Lumbsch, T.H., Grossart, H.-P., Masigol, H., Timling, I., Hiiesalu, I., Oja, J., Kupagme, J.Y., Geml, J., Alvarez-Manjarrez, J., Ilves, K., Loit, K., Adamson, K., Nara, K., Küngas, K., Rojas-Jimenez, K., Bitenieks, K., Irinyi,

L., Nagy, L.G., Soonvald, L., Zhou, L.-W., Wagner, L., Aime, M.C., Öpik, M., Mujica, M.I., Metsoja, M., Ryberg, M., Vasar, M., Murata, M., Nelsen, M.P., Cleary, M., Samarakoon, M.C., Doilom, M., Bahram, M., Hagh-Doust, N., Dulya, O., Johnston, P., Kohout, P., Chen, Q., Tian, Q., Nandi, R., Amiri, R., Perera, R.H., dos Santos Chikowski, R., Mendes-Alvarenga, R.L., Garibay-Orijel, R., Gielen, R., Phookamsak, R., Jayawardena, R.S., Rahimlou, S., Karunarathna, S.C., Tibpromma, S., Brown, S.P., Sepp, S.-K., Mundra, S., Luo, Z.-H., Bose, T., Vahter, T., Netherway, T., Yang, T., May, T., Varga, T., Li, W., Coimbra, V.R.M., de Oliveira, V.R.T., de Lima, V.X., Mikryukov, V.S., Lu, Y., Matsuda, Y., Miyamoto, Y., Kõljalg, U., Tedersoo, L., 2020. FungalTraits: a user-friendly traits database of fungi and fungus-like stramenopiles. Fungal Diversity 105, 1–16. https://doi.org/10.1007/s13225-020-00466-2

Rosling, A., Cox, F., Cruz-Martinez, K., Ihrmark, K., Grelet, G.-A., Lindahl, B.D., Menkis, A., James, T.Y., 2011. Archaeorhizomycetes: Unearthing an Ancient Class of Ubiquitous Soil Fungi. Science 333, 876–879. https://doi.org/10.1126/science.1206958

Tedersoo, L., Bahram, M., Põlme, S., Kõljalg, U., Yorou, N.S., Wijesundera, R., Ruiz, L.V., Vasco-Palacios, A.M., Thu, P.Q., Suija, A., Smith, M.E., Sharp, C., Saluveer, E., Saitta, A., Rosas, M., Riit, T., Ratkowsky, D., Pritsch, K., Põldmaa, K., Piepenbring, M., Phosri, C., Peterson, M., Parts, K., Pärtel, K., Otsing, E., Nouhra, E., Njouonkou, A.L., Nilsson, R.H., Morgado, L.N., Mayor, J., May, T.W., Majuakim, L., Lodge, D.J., Lee, S.S., Larsson, K.-H., Kohout, P., Hosaka, K., Hiiesalu, I., Henkel, T.W., Harend, H., Guo, L., Greslebin, A., Grelet, G., Geml, J., Gates, G., Dunstan, W., Dunk, C., Drenkhan, R., Dearnaley, J., De Kesel, A., Dang, T., Chen, X., Buegger, F., Brearley, F.Q., Bonito, G., Anslan, S., Abell, S., Abarenkov, K., 2014. Global diversity and geography of soil fungi. Science 346, 1256688. https://doi.org/10.1126/science.1256688
